# Membrane Lipid-K_IR_2.x Channel Interactions Enable Hemodynamic Sensing in Cerebral Arteries

**DOI:** 10.1101/455410

**Authors:** Maria Sancho, Sergio Fabris, Bjorn O. Hald, Shaun L. Sandow, Tamie L. Poepping, Donald G. Welsh

## Abstract

Inward rectifying (K_IR_) K^+^ channels are present in cerebral arterial smooth muscle and endothelial cells, a tandem arrangement suggestive of a dynamic yet undiscovered role for this channel. We explored whether vascular K_IR_ channels were uniquely modulated by membrane lipids and hemodynamic stimuli. A K_IR_ current was isolated in smooth muscle and endothelial cells of rat cerebral arteries and molecular analyses confirmed K_IR_2.1/K_IR_2.2 mRNA and protein expression. Electrophysiology next revealed that endothelial K_IR_ was sensitive to phosphatidylinositol 4,5- bisphosphate (PIP_2_), with depletion impairing flow-induced activation of the channel. In contrast, smooth muscle K_IR_ was sensitive to membrane cholesterol, with sequestration blocking pressure’s ability to inhibit this channel. Membrane lipids helped confer K_IR_ mechanosensitivity to intact arteries; virtual models then reconceptualised K_IR_ as a dynamic regulator of basal tone development. We conclude that specific membrane lipid-K_IR_ interactions enable unique channel populations to sense hemodynamic stimuli and set brain perfusion.

## Introduction

Cerebral perfusion is controlled by a complex network of resistance arteries (Segal and Duling, 1986; Segal, 2000) responsive to stimuli derived from metabolism (Filosa et al., 2006), neural activity (Si et al., 2002; Shih et al., 2012), and hemodynamic forces (Knot and Nelson, 1998; Drouin and Thorin, 2009). Irrespective of their origin, vasoactive stimuli set arterial tone through receptors and linked transduction pathways that alter smooth muscle [Ca^2+]^ and the sensitivity of the contractile apparatus to this divalent cation. The concentration of cytosolic Ca^2+^ is in turn set by membrane potential (V_M_) and the graded opening of voltage-gated Ca^2+^ channels (Welsh et al., 2000; Welsh et al., 2002).

Arterial VM reflects a dynamic interplay among depolarizing inward- and hyperpolarizing outward currents, the latter enabled by voltage-gated, Ca^2+^-activated, ATP-sensitive and inward rectifying (K_IR_) K^+^ channels (Nelson and Quayle, 1995; Jackson, 2005). An intriguing feature of vascular ion channels is that they are typically expressed either in smooth muscle or the endothelium (Jackson, 2005). While distinct in their expression pattern, all impact arterial V_M_ as vascular cells are electrically coupled to one another (Diep et al., 2005; Tran et al., 2012). K_IR_ channels break from convention in the cerebral circulation as recent work has noted robust currents in both cell types (Wu et al., 2007; Kochukov et al., 2014; Sancho et al., 2017; Longden et al., 2017).This observation implies the vascular K_IR_ channels may be more than a resting conductance to passively set tone (Nelson and Quayle, 1995; Standen and Quayle, 1998).

Vascular K_IR_ channels are composed of 4 α-subunits from the K_IR_2.x subfamily whose electrophysiological profile includes strong inward rectification, a small outward hump, extracellular K^+^ potentiation and voltage-dependent blockade by micromolar Ba^2+^ (Wu et al., 2007; Bradley et al., 1999; Schram et al., 2003). While K_IR_2.x channels are rarely viewed as a phosphorylation target, previous studies in culture systems revealed the channel interact with the surrounding lipid membrane microenvironment (Huang et al., 1998; Rohacs et al., 2003; Hilgerman, 2007). Of note are biophysical studies that have characterized the ability of phosphatidylinositol 4,5-bisphosphate (PIP_2_) and cholesterol to facilitate and suppress K_IR_2.x activity, respectively (Levitan, 2009; Rosenhouse-Dantsker et al., 2014). Although intriguing, it remains unclear whether such lipid-channel interactions influence arterial tone and if so what stimuli utilize this unique form of regulation. One possibility, gleaned from cell model systems and based upon the presumptive mechanosensitivity of K_IR_ channels, is that specific membrane lipid interactions help confer hemodynamic sensing and setting of basal V_M_ and tone in the cerebral circulation (Wu et al., 2007; Ahn et al., 2017).

This study sought to redefine K_IR_ channels in the cerebral circulation, examine their regulation by membrane lipids and determine whether these unconventional signaling molecules contribute to hemodynamic sensing. Experiments performed on freshly isolated vascular cells and intact cerebral arteries, incorporated the use of patch clamp electrophysiology, vessel myography, qPCR, immunolabeling, and computer modeling. Work first confirmed the existence of a smooth muscle and endothelial K_IR_ channel pool composed of K_IR_2.1 and K_IR_2.2 subunits. Each K_IR_ channel pool was uniquely regulated by membrane lipids, including PIP_2_, which augmented endothelial K_IR_ activity while cholesterol suppressed smooth muscle K_IR_. Further experimentation revealed that intravascular pressure diminished smooth muscle K_IR_ activity in a cholesterol dependent manner whereas the flow sensitivity of endothelial K_IR_ was tied to PIP_2_ content. Incorporating these observations, computational modeling re-envisioned how K_IR_ channels dynamically interact with other conductances setting basal V_M_ and tone development in the cerebral circulation. This study is the first to illustrate that specific membrane lipid-K_IR_ interactions enable particular populations of K_IR_ channels to sense hemodynamic stimuli and set the basis of tone development and blood flow delivery in the cerebral circulation.

## Results

### K_IR_ Channel Expression in Cerebral Vascular Cells

This examination began with a characterization of K_IR_2.x expression in cerebral arterial smooth muscle and endothelium. Using patch-clamp electrophysiology and freshly isolated cells, whole cell K_IR_ currents (ramps, -100 to +20 mV) were monitored in a 60 mM K^+^ solution as Ba^2+^ (1-100 µM), a selective K_IR_ inhibitor, was sequentially increased. Figures 1A & 1B denotes the presence of a Ba^2+^-sensitive current in both cell types at voltages negative to the K^+^ equilibrium potential (E_*K*_). These currents displayed micromolar sensitivity to Ba^2+^ (Figure 1C), with the amplitude of endothelial K_IR_ (-20.04±1.32 pA/pF) exceeding that of smooth muscle (-1.73±0.07 pA/pF) by ~12 fold (-100 mV). The IC_50_ of Ba^2+^ block (-60 mV) for the smooth muscle and endothelial current was 1.05±0.02 µM and 3.5±0.5 µM, respectively (Figure 1D). Quantitative PCR analysis of arteries and isolated smooth muscle/endothelial cells revealed mRNA expression of K_IR_2.1, K_IR_2.2 and K_IR_2.4 (Figure 1E). Complementary western blot analysis confirmed protein expression of K_IR_2.1/ K_IR_2.2 in whole arteries (Figure 1F); immunohistochemical analysis revealed that both subunits were present in smooth muscle and endothelium near cell borders; no positive labeling was observed for K_IR_2.4 (Figure 1G).

**Figure 1.**
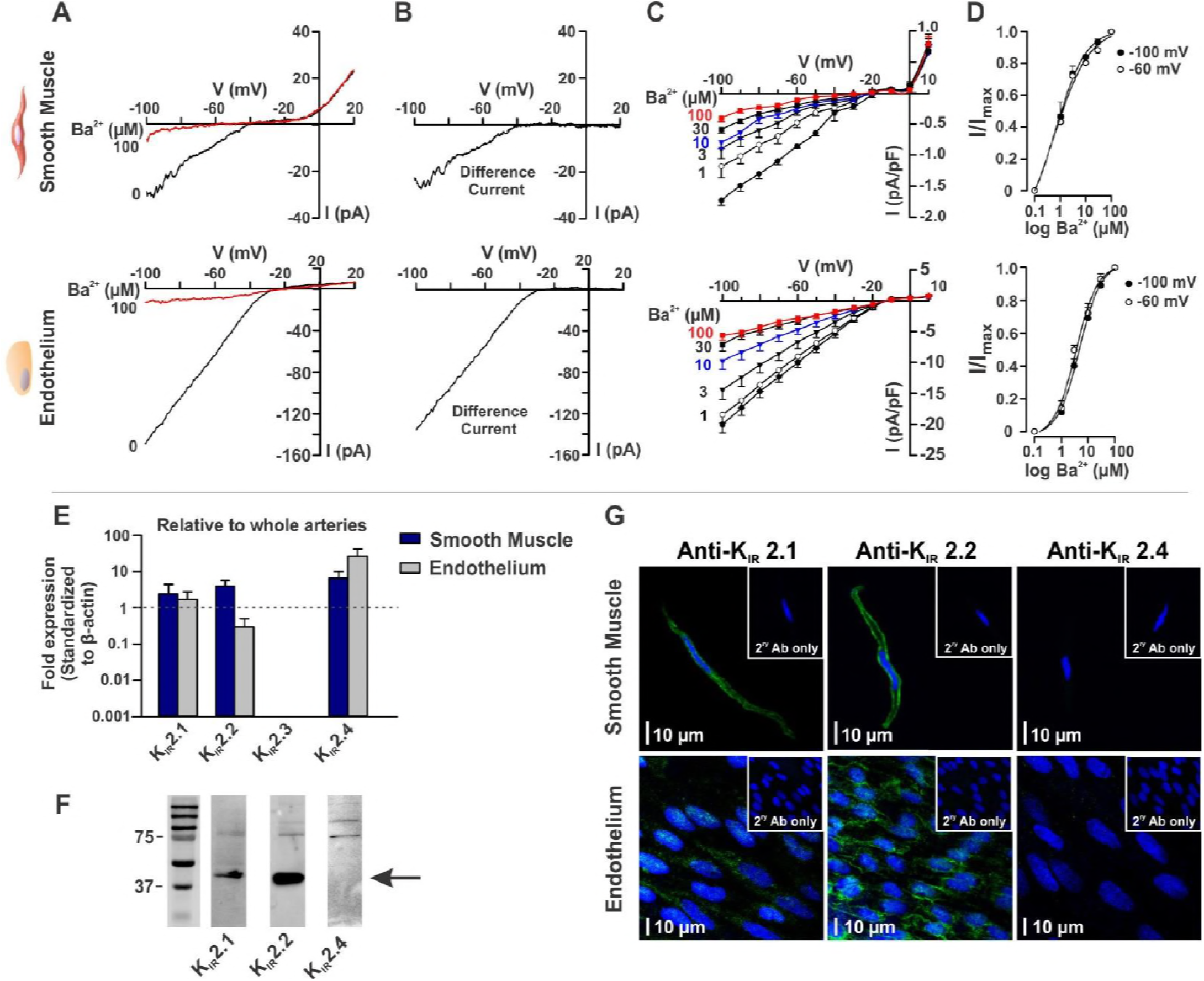
K_IR_2.x channels are present in rat cerebral arterial smooth muscle and endothelial cells. Whole-cell K^+^ currents were measured in isolated cells bathed in 60 mM K^+^ using a voltage ramp protocol (-100 to +20 mV) prior to and following the addition of BaCl_2_ (K_IR_ blocker; 1 to 100 μM). **A & B**, Representative whole-cell current recordings and Ba^2+^-subtracted currents in freshly isolated smooth muscle (top) and endothelial (bottom) cells. **C**, Summary plots (n=9 cells) illustrate the effect of Ba^2+^ on whole-cell currents. **D**, Ba^2+^ induces a concentration-dependent blockade of K_IR_ current at both -100 mV and -60 mV (n=9). IC50 values (at -60 mV) for the smooth muscle and endothelial K_IR_ were 1.05±0.02 µM and 3.5±0.5 µM, respectively. Data are means±SE. **E**, qPCR analysis of isolated smooth muscle and endothelial cells highlights the presence of K_IR_ 2.1, K_IR_2.2 and K_IR_ 2.4 mRNA. Data was derived from 6 experiments performed on 6 separate animals were expressed relative to β-actin. **F**, Western blot analysis of whole cerebral arteries further confirm K_IR_ 2.1 and K_IR_ 2.2 protein expression (n=4 experiments from 4 animals). **G**, Immunohistochemistry of isolated smooth muscle cells (top) and the intact endothelial layer (bottom) reveal protein expression of K_IR_ 2.1/K_IR_2.2 (green) but not K_IR_2.4 subunits. Nuclei were stained with DAPI (blue); and assay controls were performed with no primary antibodies. Each experiment was tested on cells for 4 different animals; photomicrographs are representative of 15-20 cells per animal.

### Dependence of Vascular K_IR_ Channels on Membrane PIP_2_

Patch-clamp experiments next explored the sensitivity of each pool of K_IR_2.x channels to membrane PIP_2_ manipulation. Upon membrane rupture, whole-cell currents were continuously monitored to assess time-dependent changes in response to agents dissolved in the patch pipette. Figure 2A highlights that irrespective of the cell type, K_IR_ activity was stable over time when dialyzed with a standard concentration of ATP (2.5 mM). Dropping the ATP concentration to zero blocks PI4 and PI5 kinases (PI4K & PI5K) and induced a marked run down of the endothelial (~39%) but not the smooth muscle K_IR_ current over 12 min (Figure 2B). The steady decrease in the endothelial K_IR_ current was prevented by adding the water-soluble short-chain PIP_2_ analog (dioctanoyl-PIP_2_, diC8-PIP_2_, 50 μM) to the pipette, a finding consistent with PIP_2_ gating this channel pool. In contrast, dialysis of endothelial cells with 5 mM ATP, which augmented the K_IR_ current by ~38%, a response blocked by neomycin, an aminoglycoside that binds PIP_2_ impairing electrostatic interaction between this lipid and K_IR_ channels (Figure 2C). The smooth muscle K_IR_ current was unaffected by elevated ATP (5 mM), a finding consistent with PIP_2_ being a minor regulator of this particular channel pool.

**Figure 2.**
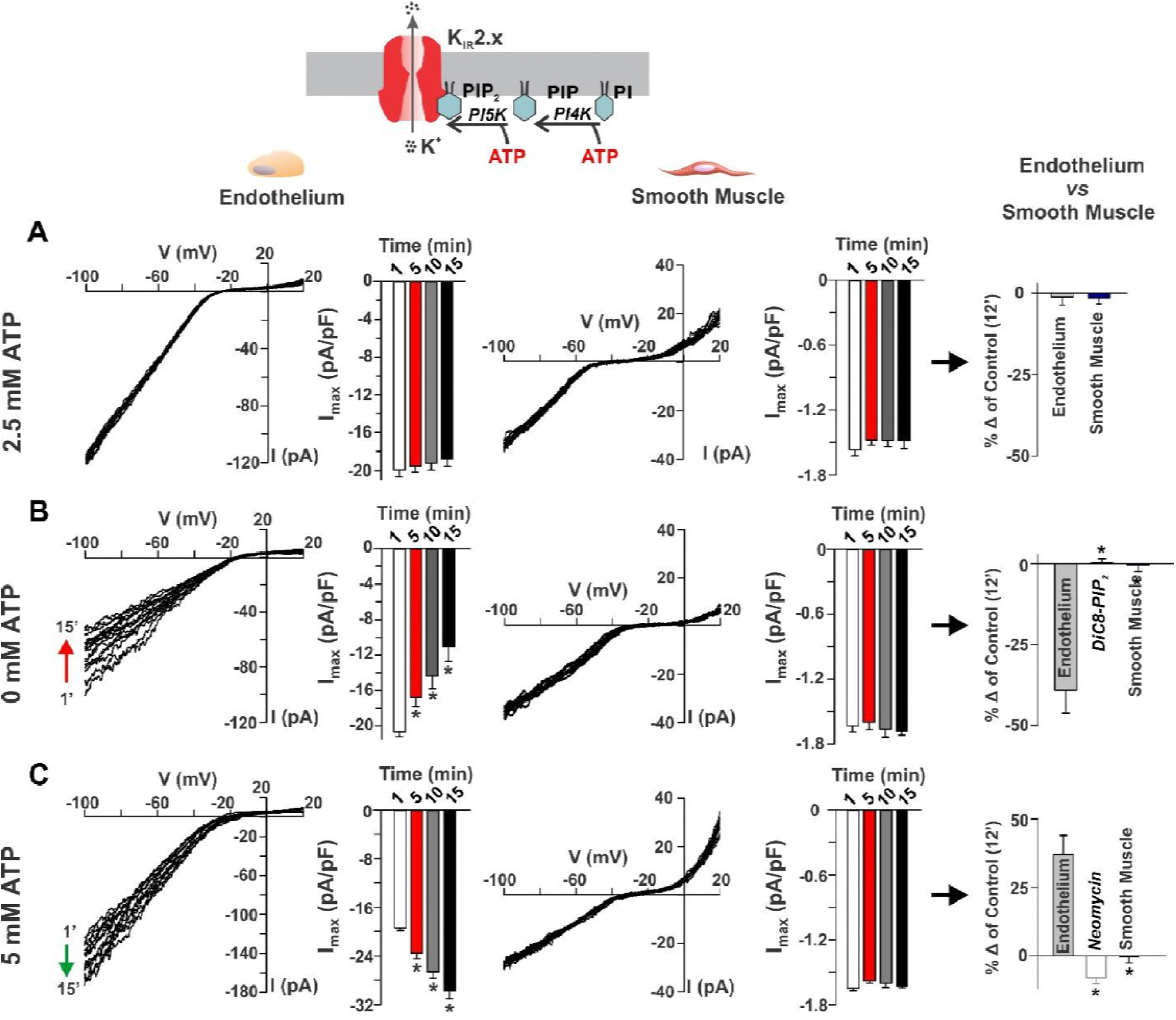
Intracellular [ATP] alters K_IR_ channel activity in cerebral arterial endothelial but not smooth muscle cells. Top: schematic diagram highlights importance of ATP in driving PIP_2_ synthesis via PI4K/PI5K. Representative traces and summary data of peak inward current (-100 mV; 60 mM K^+^ in the bath) in freshly isolated endothelial (left) and smooth muscle cells (middle) over a 15 min dialysis period. The recording pipette retained 2.5 mM ATP (**A;** n=8, RM-ANOVA, Tukey post-hoc analysis), 0 ATP ± 50 µM DiC8-PIP_2_ (**B**; n=6, RM-ANOVA, Tukey post-hoc analysis) or 5 mM ATP ± 50 µM neomycin (**C**; n=6, RM-ANOVA, Tukey post-hoc analysis). Summary data (n= 6-8; unpaired *t* test; right) was expressed as percent effect (K_IR_ current at 12 min/0 min). Data are means±SE and * denotes significant difference at *P* ≤ 0.05.

A broader range of agents was subsequently applied to examine the regulatory impact of PIP_2_ on vascular K_IR_ channels. Disrupting PIP_2_-protein interactions with neomycin (50 μM) supressed inward K_IR_ current by ~42% (measured at -100 mV); outward current (+20 mV) reflecting non K_IR_ currents was unaffected (Figures 3A & 3C; Figure 3- figure supplement 1). Dialyzing cells with the water-soluble PIP_2_ analog diC8-PIP_2_ (50 μM) plus 2.5 mM ATP enhanced the inward K_IR_ current (~47%) while having no impact on outward current (Figure 3- figure supplement 1). This potentiation was minimized by adding neomycin to the pipette solution (Figure 3C, bottom). Similar treatment of smooth muscle cells failed to alter the inward or the outward component of the whole cell current, the latter dominated by voltage-gated and large conductance Ca^2+^ activated K^+^ channels (Figures 3B & 3C). Blocking PI-4K with wortmannin (50 μM) or adenosine (2 mM) in the pipette induced a selective time-dependent rundown of inward (K_IR_, ~30-35%; Figures 3D & 3F) but not outward current in cerebral arterial endothelial cells (Figure 3- figure supplement 1). Parallel experiments in smooth muscle cells revealed no change in whole cell current over time (Figures 3E & 3F). We conclude that endothelial but not smooth muscle K_IR_ channels are sensitive to membrane PIP_2_ (Figure 3- figure supplement 2), a lipid that promotes channel stabilization in a preferred open state.

**Figure 3.**
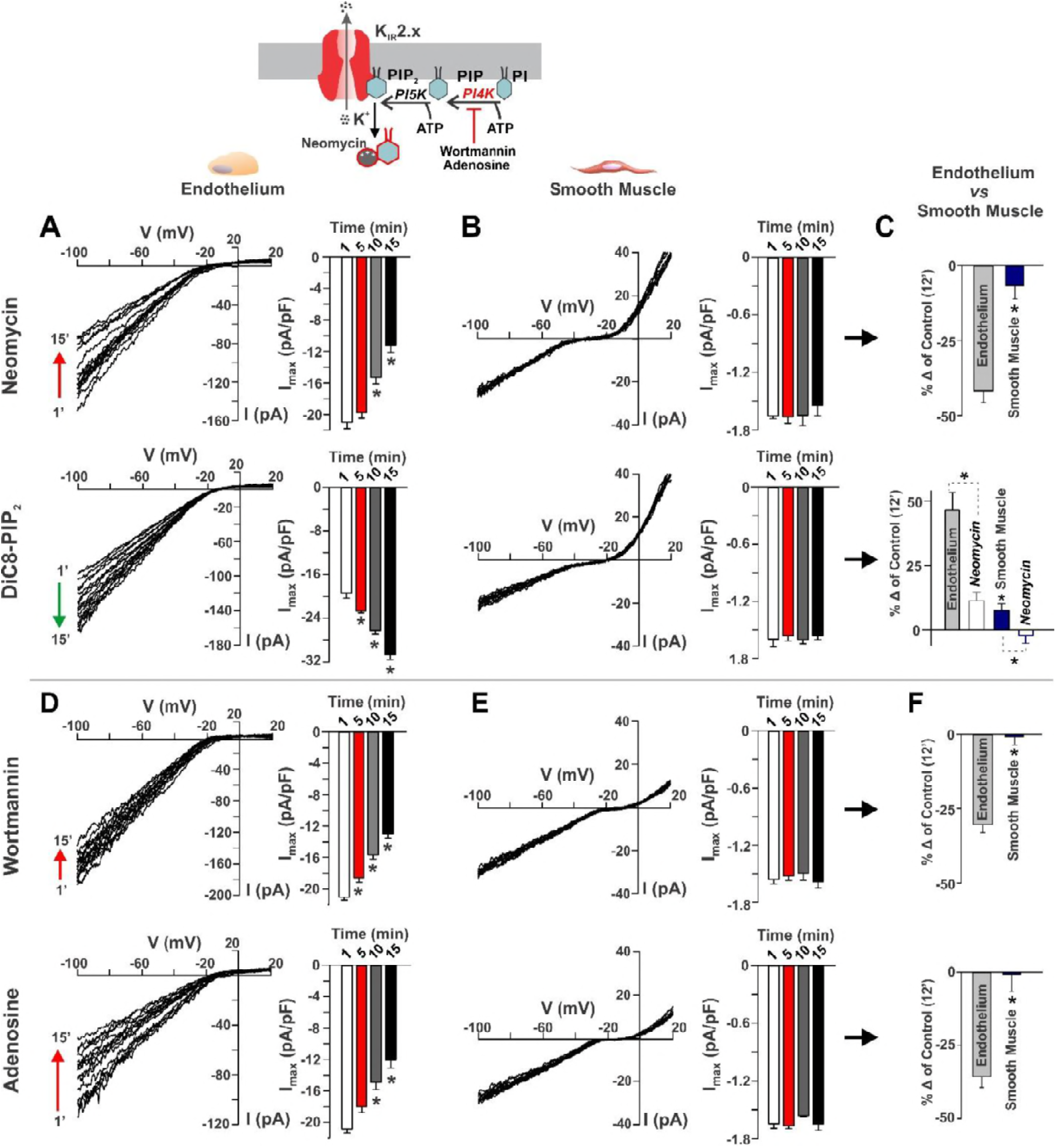
PIP_2_ drives cerebral arterial endothelial but not smooth muscle K_IR_ channel activity. Top: schematic diagram highlighting the pharmacological manipulations used to alter membrane PIP_2_ content. Representative recordings and summary data of peak inward current (- 100 mV; 60 mM K^+^ in the bath) in native endothelial (**A** & **D**) and smooth muscle cells (**B** & **E**) during the dialysis (15 min) of neomycin (50 µM; n=9; RM-ANOVA, Tukey post hoc analysis), DiC8-PIP_2_ ± neomycin (50 µM; n=8, RM-ANOVA, Tukey post hoc analysis), wortmannin (50 µM; n=6, RM-ANOVA, Tukey post hoc analysis), or adenosine (2 mM; n=7, RM-ANOVA, Tukey post hoc analysis). **C** & **F**, Summary data (n= 6-8; unpaired *t* test; right) was expressed as percent effect (K_IR_ current at 12 min/0 min). Data are means±SE and * denotes significant difference at *P* ≤ 0.05.

### Cholesterol Sensitivity of Vascular K_IR_ Channels

As K_IR_ channels are sensitive to membrane cholesterol content (Levitan, 2009), experiments were performed to assess the impact of this lipid on vascular currents. Depleting cells of cholesterol (5 mM methyl-β-cyclodextrin, MβCD), increased K_IR_ activity at -100 mV, an effect that was particularly prominent in smooth muscle (~100%) rather than endothelial cells (-30%; Figure 4A). This time-dependent potentiation was abrogated by dialyzing both cell types with water-soluble cholesterol (5 mM) combined with MβCD (~10% change; Figure 4B); outward current at +20 mV, representative of no K_IR_ channel activity was unaffected (Figure 3- figure supplement 1). These findings collectively reveal that cholesterol is a dominant lipid regulator of smooth muscle K_IR_, promoting channel stabilization in a preferred silent state (Figure 3- figure supplement 2).

**Figure 4.**
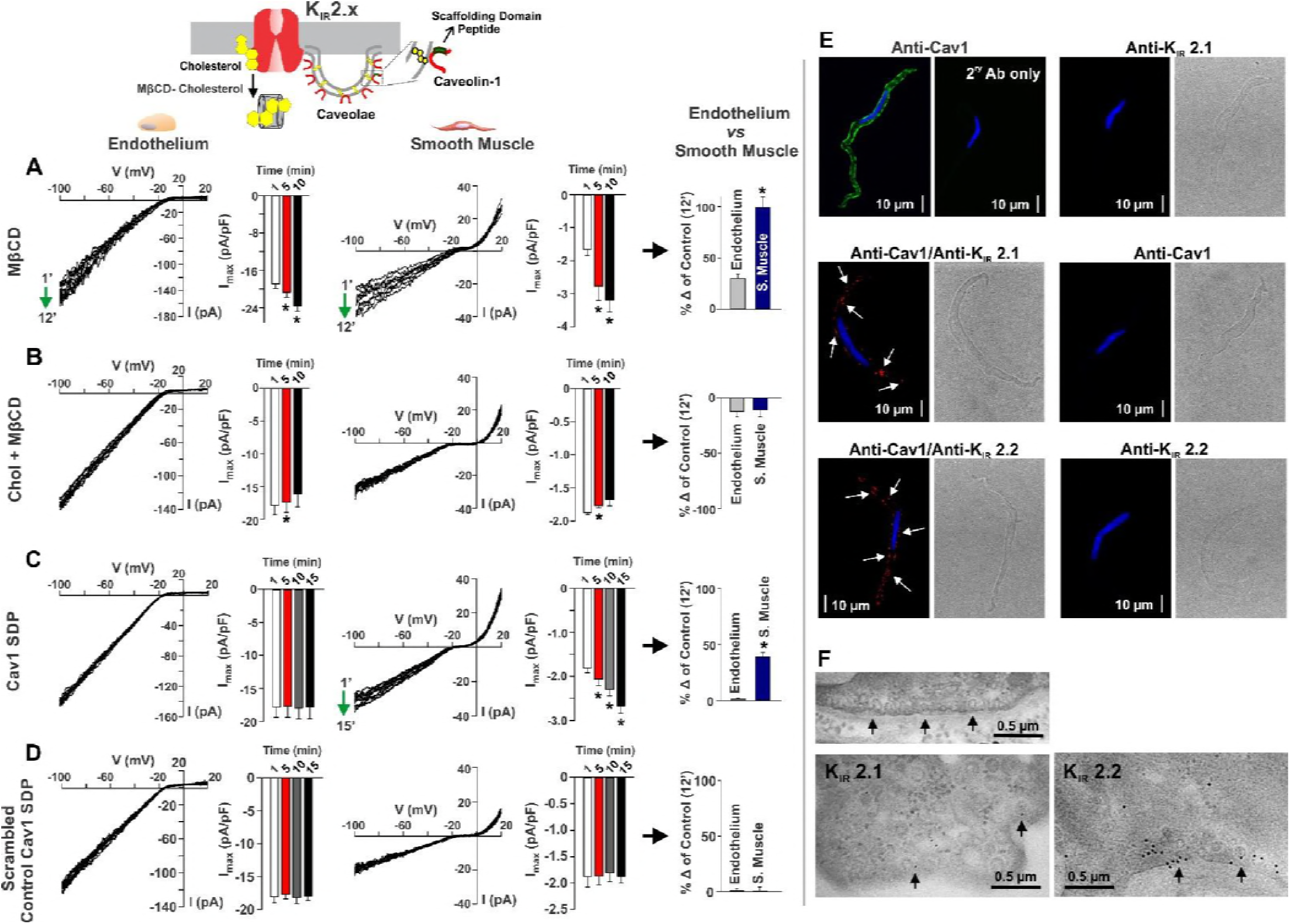
Cholesterol is a key lipid regulator of cerebral arterial smooth muscle but not endothelial K_IR_ channels. Top: schematic diagram highlighting the pharmacological manipulations used to modify membrane cholesterol content. Representative traces and summary data of peak inward current (-100 mV; 60 mM K^+^ in the bath) in isolated endothelial and smooth muscle cells over a 15 min dialysis period. The recording pipette retained methyl-β-cyclodextrin ± cholesterol (**A** & **B**; 5 mM; n=7-8, RM-ANOVA, Tukey post hoc analysis), Cav1 SDP (**C**; 10 µM; n=6-7, RM-ANOVA, Tukey post hoc analysis), or Cav1 SDP scrambled control (**D**; 10 µM; n=6-9, RM-ANOVA, Tukey posthoc analysis). Summary data (n= 6-9; unpaired *t* test) was expressed as percent effect (K_IR_ current at 12 min/0 min). Data are means±SE and * denotes significant difference at *P* ≤ 0.05. **E**, The performance of immunohistochemistry (left panel) and the proximity ligation assay on isolated smooth muscle cells revealed the presence and close apposition (< 40 nm) of K_IR_ 2.1/K_IR_ 2.2 with Cav1. Nuclei were stained with DAPI (blue); controls were performed with one or both primary antibodies removed. Each experiment was tested on cells from 4 different animals; photomicrographs are representative of 15-20 cells per animal. **F**, Transmission electron microscopy and immunogold labeling of K_IR_2.1 and K_IR_2.2 in rat cerebral arteries. K_IR_2.1 (left, bottom) and K_IR_2.2 (right, bottom) labeling (arrowheads) was confined to the plasma membrane in association with caveolae; further shown in control unlabelled sections (top). Each photomicrograph is representative of 4 independent preparations.

Since ion channels often localize to caveolae, structures enriched with cholesterol, a subset of experiments examined whether caveolin-1 (Cav1), the dominant vascular isoform, modulated K_IR_ channels. K_IR_ activity rose ~40% in smooth muscle cells dialyzed with solution containing Cav1 SDP (10 μM), a scaffolding domain peptide that disrupts Cav1 signaling; endothelial K_IR_ was largely unaffected (Figure 4C). Control experiments with the scrambled peptide had no impact on basal currents in either cell type (Figure 4D). Immunohistochemistry confirmed Cav1 expression in isolated smooth muscle cells, particularly near cell borders (Figure 4E, left panel). A proximity ligation assay revealed that Cav1 and K_IR_2.1/ K_IR_2.2 reside within 40 nm of one another as punctuate red fluorescent product was detected in smooth muscle cells. Fluorescent product was absent in control experiments when one or both primary antibodies were omitted (Figure 4E). Immunogold labeling subsequently confirmed that both K_IR_2.1 and K_IR_ 2.2 were limited to the smooth muscle plasma membrane at or near caveolae (Figure 4F).

### Mechanosensitivity of Vascular K_IR_ Channels

To test whether K_IR_ channels were sensitive to hemodynamic forces, whole cell currents were monitored in vascular cells (bathed with 20 mM K^+^) in response to perturbations that simulate intravascular pressure (cell swelling and pipet pressurization) or flow. Reducing bath osmolarity to induce cell swelling suppressed the smooth muscle K_IR_ current (~30% after 6 min, -100 mV) whereas negative pipette pressurization induced an immediate rise (~41% at -45mmHg; Figures 5A & 5B). The latter manipulation is challenging given its potential to change seal resistance. Neither pressure perturbation impacted the endothelial K_IR_ current (Figures 5A & 5B) or outward K^+^ current in smooth muscle at +20 mV (Figure 3- figure supplement 1). Using Ba^2+^ reactivity as an index of K_IR_ activity, the impact of this divalent on denuded arteries at low (15 mmHg) and high (80 mmHg) pressure was next assessed. At low pressure, where myogenic tone is modest and vessels hyperpolarized, Ba^2+^ (30 µM) induced a robust constriction (23.7±2.8 µm); this response was attenuated (9.1±1.6 µm) at high pressure in depolarized, myogenically active vessels (Figure 5C). Barium reactivity was absent in denuded arteries from the mesenteric circulation, vessels where K_IR_ channels aren’t expressed in the smooth muscle layer (Figure 5- figure supplement 1) (Smith et al., 2008).

**Figure 5.**
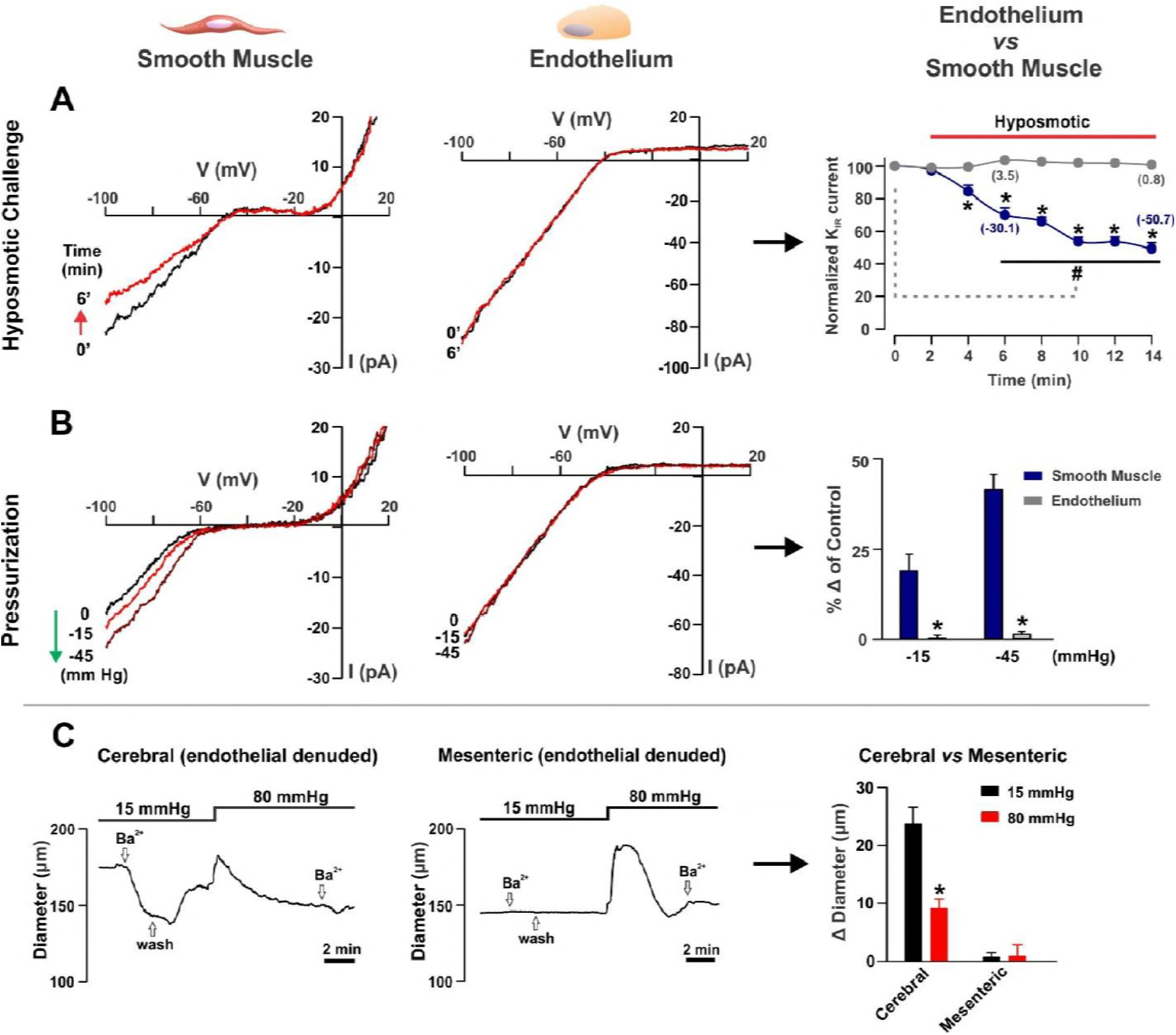
Simulated pressure impacts K_IR_ channel activity in cerebral arterial smooth muscle but not endothelium. Whole-cell currents were measured in isolated cells bathed in 20 mM K^+^ solution (voltage ramp protocol, -100 to +20 mV) prior to and following pressure simulation. **A** & **B**, Representative traces and summary plots of smooth muscle and endothelial K_IR_ channel activity (-100 mV) prior to and following a hyposmotic challenge (reduced bath osmolarity from 300 to 205 mosM; n=6, RM-ANOVA and Tukey post hoc analysis for time course data; unpaired *t* test compared the two groups at each time point) or pipette pressurization (-15 or -45 mmHg; n=7 unpaired *t* test). **C**, Representative traces and summary data highlighting the effect of Ba^2+^ (30 µM) on cerebral and mesenteric arteries pressurized to 15 or 80 mmHg (n=6, paired *t* test). Data are means±SE; *, # denotes significant difference at *P* ≤ 0.05.

As membrane cholesterol is a dominant regulator of smooth muscle K_IR_, we next assessed whether this lipid helped confer pressure-sensitivity. Cholesterol depletion (MβCD in the pipette) prevented swelling from inhibiting K_IR_ in smooth muscle cells; Cav1 SDP also impaired this pressure-sensitive response (Figure 6A; Figure 6- figure supplement 1). In contrast, PIP_2_ sequestration (neomycin in the pipette) had no effect on swelling-induced K_IR_ suppression (Figure 6B). These functional findings imply that if cholesterol is depleted in an intact artery, Ba^2+^-induced constriction (an index of smooth muscle K_IR_ activity) should rise at elevated intravascular pressure (80 mmHg). Indeed, Ba^2+^-induced constriction rose ~3 fold in denuded arteries pretreated with 5 mM MβCD (30 min) compared to control tissues (Figure 6C).

**Figure 6.**
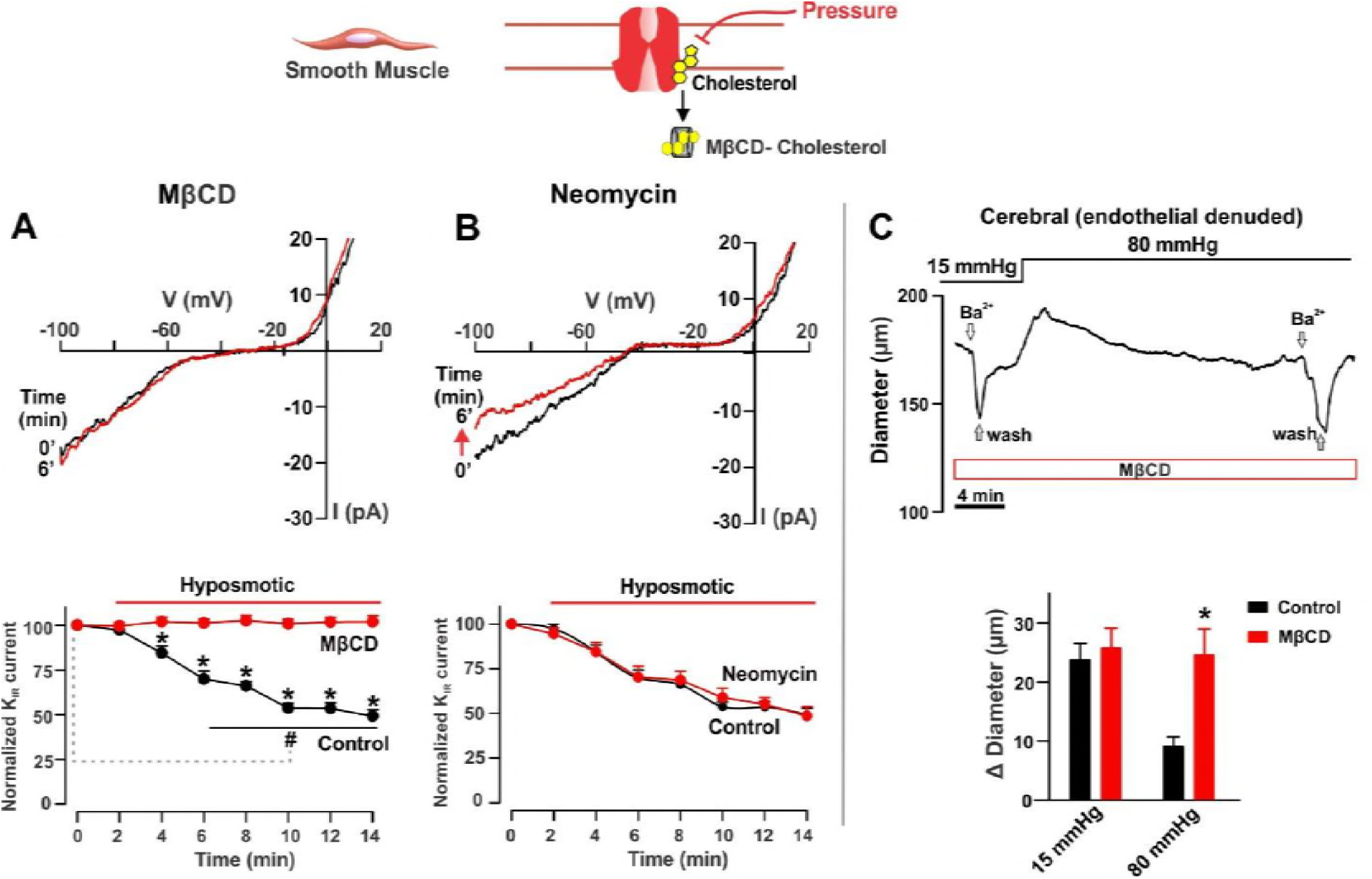
Cholesterol depletion prevents pressure from suppressing smooth muscle K_IR_ channel activity. **A & B**, Representative traces and summary plots (n=6-7; RM-ANOVA and Tukey post hoc analysis for time-course differences; unpaired *t* test for percent change in peak inward current among treatments) of smooth muscle K_IR_ channel activity (20 mM K^+^ in the bath; voltage ramp, -100 to +20 mV), prior to and following hyposmotic challenge. Pipette solution contained methyl-β-cyclodextrin (**A**; 5 mM) or neomycin (**B**; 50 µM). **C**, Representative trace and summary plot highlighting the effect of Ba^2+^ (30 µM) on methyl-β-cyclodextrin treated cerebral arteries (5 mM, 30 min) pressurized to 15 or 80 mmHg (n=6, unpaired *t* test). Cerebral arteries were denuded of endothelium using air bubble treatment. Data are means±SE and *, # denotes significant difference at *P* ≤ 0.05.

Patch-clamp experiments next ascertained the flow sensitivity of cerebral vascular K_IR_ channels by monitoring whole cell currents in a custom flow chamber (laminar shear stress ~10 mPa). Figure 7A notes that elevated flow increased the endothelial K_IR_ current (~30% at -100 mV) without modifying whole-cell outward current at +20 mV (Figure 3- figure supplement 1). In contrast, smooth muscle K_IR_ activity (measured at -100 mV) was unaffected by this stimulus. Since PIP_2_ is a main lipid regulator of endothelial K_IR_ channels, we then explored whether this phospholipid fostered flow activation. As expected, PIP_2_ sequestration (neomycin in the pipette) eliminated flow activation of endothelial K_IR_ while cholesterol depletion (MβCD in the pipette) had no effect (Figure 7B). These observations imply that endothelial K_IR_ contributes to flow-mediated vasodilation, and that membrane PIP_2_ helps confer this response by sustaining the channel in an open state. Consistent with this hypothesis, flow-induced dilation (0-6 µl/min, 2 µl/min steps) was substantively impaired by Ba^2+^ (30 µM; intra- or extra-luminal application) and neomycin (50 µM, intraluminal application) (Figure 7C; Figure 7- figure supplement 1).

**Figure 7.**
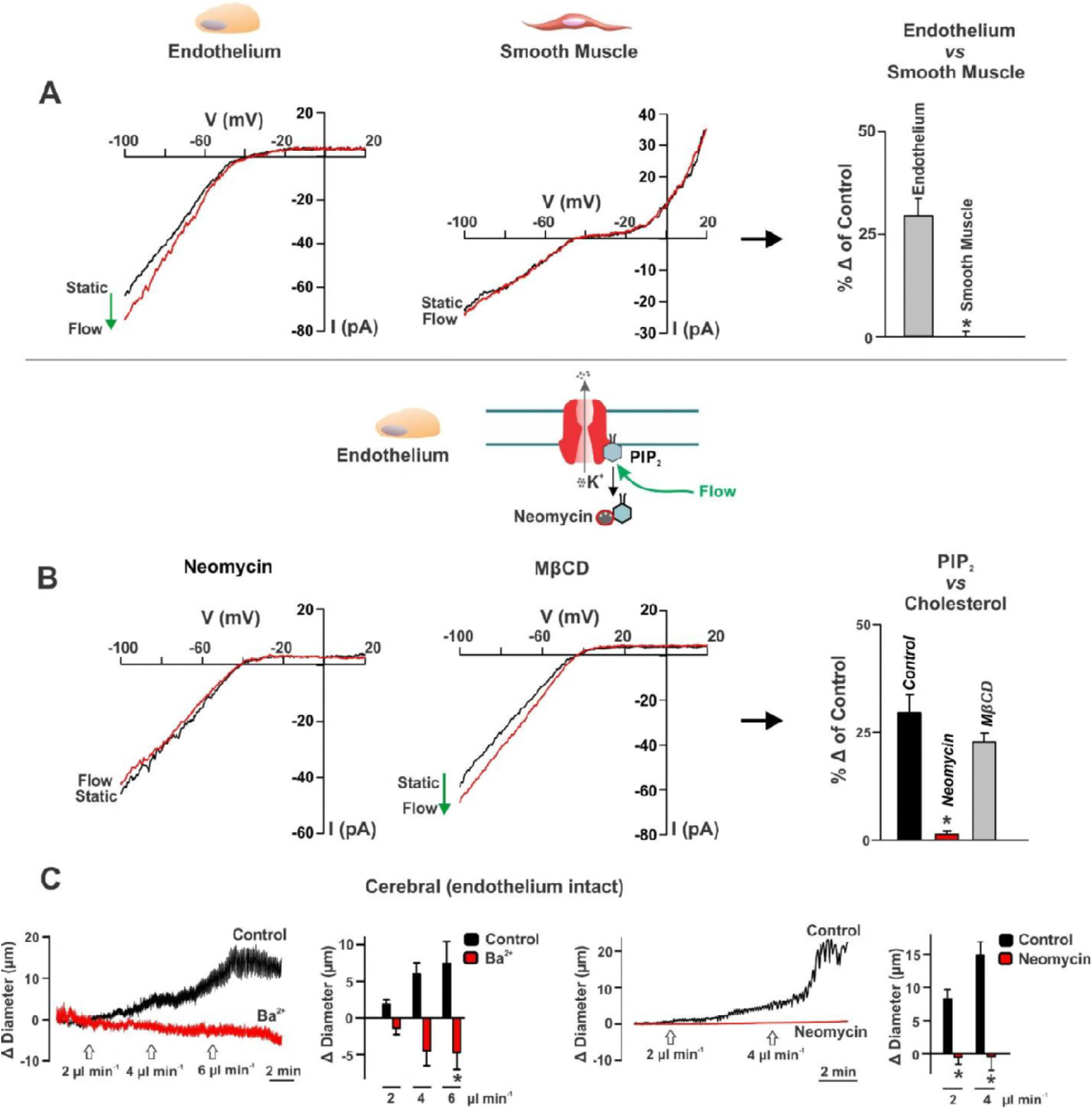
Laminar flow activates endothelial but not smooth muscle K_IR_ channels, an effect impaired by PIP_2_ sequestration. Whole-cell currents were measured in cerebral arterial cells bathed in 20 mM K^+^ solution (voltage ramp, -100 to +20 mV), prior to and after the induction of shear stress (~10 mPA). **A**, Representative traces and summary data (n=6-7, unpaired *t* test) of the endothelial and smooth muscle K_IR_ current (-100 mV) in response to laminar flow. **B**, Representative traces and summary plots (n=7-8, paired *t* test) of endothelial K_IR_ channel activity exposed to laminar flow; the pipette solution contained neomycin (50 µM), or methyl-β-cyclodextrin (5 mM). **C**, Representative traces and summary data (n=6, paired *t* test) highlighting the effect of intraluminal Ba^2+^ (30 µM) or neomycin (50 µM) on flow-mediated vasodilation (0 to 4 or 6 µl/min) in endothelium-intact cerebral arteries. Data are means±SE and * denotes significant difference at *P* ≤ 0.05.

### Vascular K_IR_ Channels Interact to Control Arterial V_M_ and Tone

To conceptually explore how smooth muscle and endothelial K_IR_ channels work interactively to set arterial V_M_ and tone, two computational models were used, the first being a single virtual artery (Hald, 2016) and the second a cerebral vascular network (6 penetrating arteries interconnected with surface vessels) structured on the multi-photon observations of Shih et al., 2009 (Figure 8A). Briefly, vessel structure consisted of 1-2 layers of smooth muscle and 1 layer of endothelium, with cells arranged perpendicular and parallel to the vessel’s long axis. Cells were interconnected with linear resistors representing gap junctions and their ionic properties were modeled as two components, one reflecting K_IR_ and the second all other currents (I_PM_; Figures 8B & 8C, left). Using the single vessel model, 2D plots (Figures 8B & 8C middle) showed that modest changes in endothelial or smooth muscle K_IR_ activity dynamically altered arterial V_M_ and diameter. This balanced result arose despite K_IR_ amplitude and cell number being starkly different among the two cell types. Colored voltage maps of the virtual network better illustrated the shift in arterial V_M_ as endothelial/smooth muscle K_IR_ activity was increased and decreased by ± 25% (Figures 8B & 8C, right).

**Figure 8.**
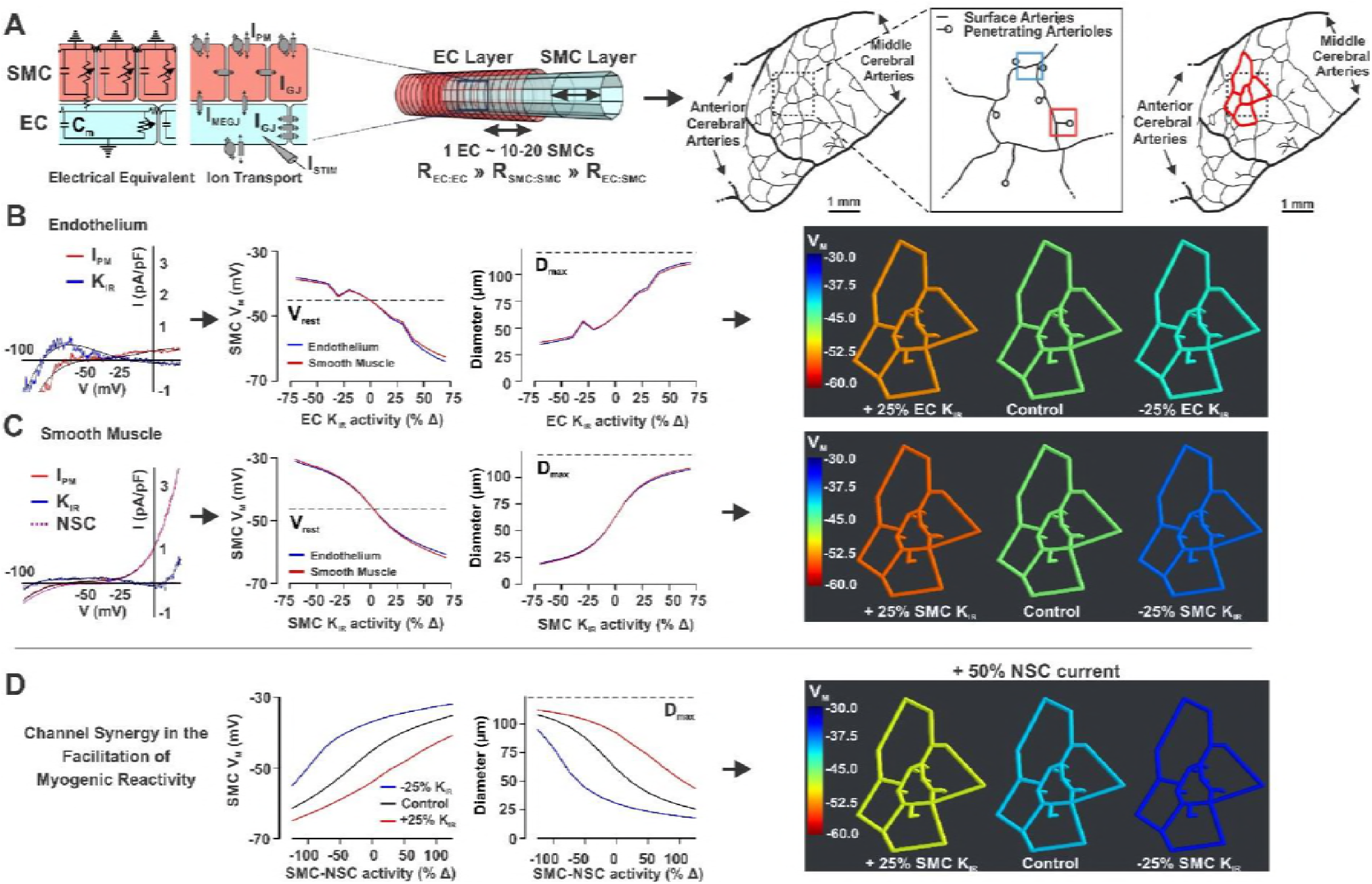
Computational modeling highlights the dynamic interplay among endothelial and smooth muscle K_IR_ channels in setting cerebral arterial V_M_ and tone. **A,** Schematic diagrams illustrating the electrical/structural design of the virtual artery (left) and network (right) used to model the influence of cerebral arterial K_IR_ channels. The virtual artery consisted of 1 layer of smooth muscle and 1 layer of endothelial, with cells interconnected with variable linear resistors to represent gap junctions. The arterial network consisted of an interconnected network of surface arteries (2 layers of smooth muscle) and six adjoining penetrating arterioles. **B & C**, The ionic properties of endothelial and smooth muscle cells was represented by a plasma membrane (IPM, red) current to which a Ba^2+^-sensitive K_IR_ component (K_IR_, blue) was added/subtracted in a graded manner (left). Modeled currents were derived from whole cell recordings collected from smooth muscle and endothelial cells bathed in physiological salt solutions. Using the single vessel model (middle), 2D plots represent the changes in arterial V_M_/diameter after systematic modification of the endothelial or smooth muscle K_IR_ activity. Colorized voltage maps (right) illustrate the shift in arterial V_M_ when altering each cellular current. D, A final subset of simulations explored the interaction among smooth muscle K_IR_ and a pressure-sensitive non-selective cation (NSC, purple in C) current. Single vessel (left) and color mapped arterial network models (right) reveal that the ability of a non-selective cation current to depolarize/constrict virtual structures was facilitated decreasing smooth muscle K_IR_ activity by 25%.

With smooth muscle K_IR_ displaying mechano-sensitivity, a final set of simulations were performed to explore how this channel population could interact with a non-selective current, itself pressure-sensitive, and known to facilitate myogenic tone (Welsh et al., 2000). Altering the activity of a pressure-sensitive cation current incorporated into the smooth muscle layer revealed that small changes in current amplitude could impact arterial V_M_ and tone (Figure 8D). Intriguingly, the magnitude of the electrical/vasomotor changes were facilitated or attenuated by increasing or decreasing smooth muscle K_IR_ activity (25%), respectively. These observations highlight the apparent synergy among pressure-sensitive channels, particularly in brain where perfusion is tightly matched with neural activity.

## Discussion

This study reconceptualised K_IR_ channels in the cerebral circulation by assessing their membrane lipid regulation and whether these molecules help confer hemodynamic sensitivity to pressure and flow. Findings revealed that channels comprised of K_IR_2.1 and K_IR_2.2 subunits are present in smooth muscle and endothelial cells, and that each cellular pool displays a unique sensitivity to membrane lipids and hemodynamic stimuli. Endothelial K_IR_ channels were particularly sensitive to PIP_2_ manipulations whereas smooth muscle K_IR_ channels were markedly altered by changes in membrane cholesterol. These lipid signaling molecules aid in conferring pressure-sensitivity to the smooth muscle K_IR_ (cholesterol dependent), and flow-sensitivity to the endothelial K_IR_ (PIP_2_ dependent), respectively. Integrating these observations into virtual models rationalized the dynamic interplay among the two K_IR_ channel pools to efficiently control voltage, cytosolic Ca^2+^ and vessel diameter over a range of potential pressures and flows. We propose that in the cerebral circulation, unique K_IR_ channel pools are required to sense hemodynamic forces and set tone required for basal perfusion control (Figure 9).

**Figure 9.**
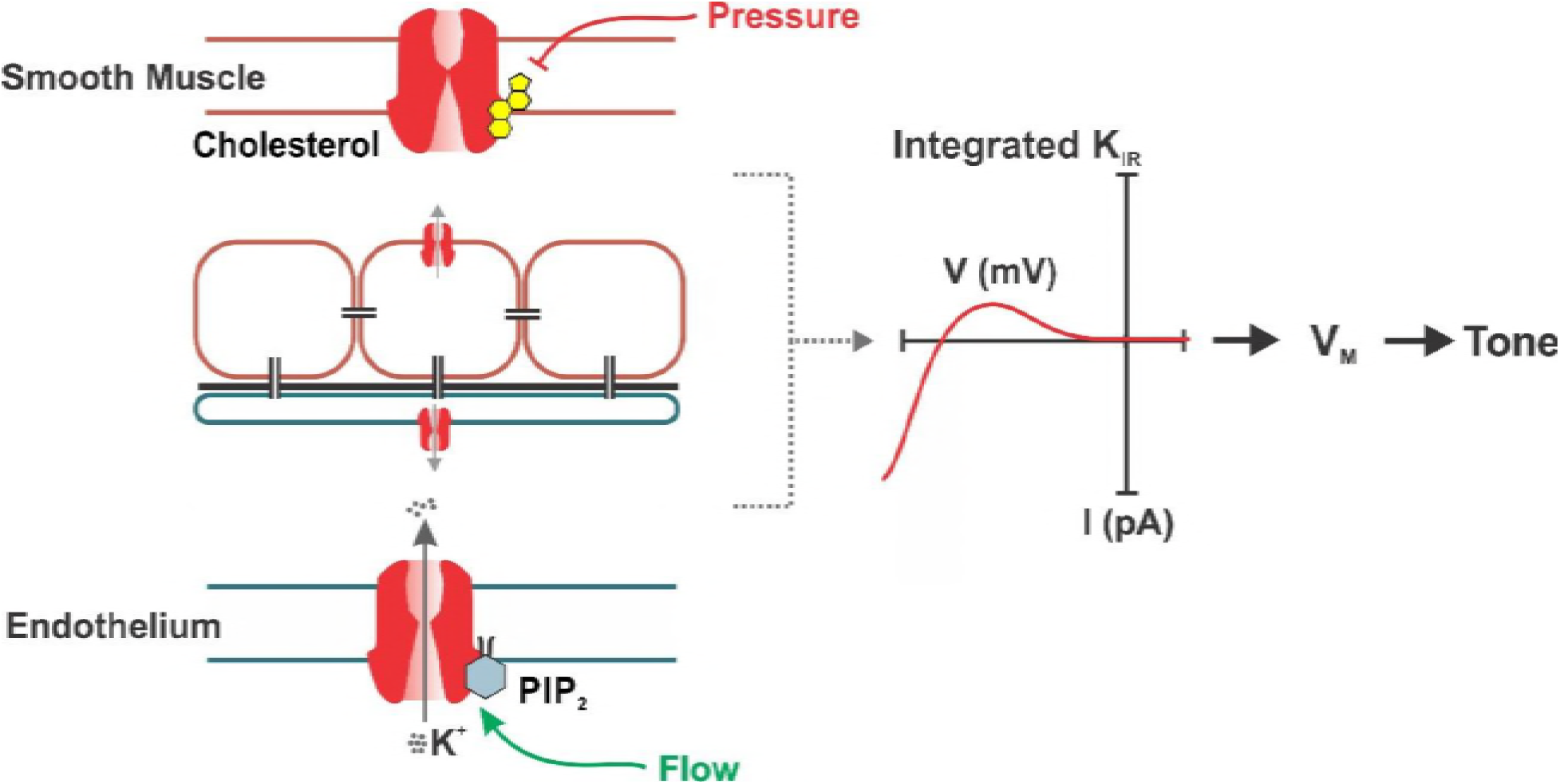
K_IR_ channels and the facilitation of hemodynamic control. Schematic diagram illustrating the unique regulation of each pool of K_IR_ channels in cerebral arterial smooth muscle and endothelium. Membrane lipids such as PIP_2_ and cholesterol are dominant regulators of K_IR_, with the former prominently activating endothelial K_IR_ while the latter preferentially inhibits smooth muscle K_IR_. These specific lipid-K_IR_ interactions help confer pressure and flow sensing, hemodynamic forces that impair and activate smooth muscle and endothelial K_IR_ channels, respectively. Both channel pools together drive an integrated K_IR_ current through which hemodynamic stimuli cooperate to control V_M_ and arterial tone.

### K_IR_2.x Channels in Cerebral Vascular Cells

Cerebral arteries form an integrated network to tune blood flow delivery in support of neural activity and brain function (Segal and Duling, 1986; Segal, 2000). Tone within this complex structure is set in a coordinated manner in response to diverse stimuli generated by tissue metabolism (Filosa et al., 2006), neuronal activity (Si et al., 2002; Shih et al., 2012), and hemodynamic forces (Knot and Nelson, 1998; Drouin and Thorin, 2009). Coordinated behaviour relies on ion channel activity and the spread of charge via gap junctions to balance arterial V_M_ and further downstream, Ca^2+^ dependent myosin light chain phosphorylation (Segal et al., 1999; Figueroa et al., 2004). As a rule, ion channels setting arterial V_M_ have distinct distribution, being present in smooth muscle or the endothelium (Jackson, 2005; Quayle et al., 1997; Thorneloe and Nelson, 2005; Ledoux et al., 2008; McNeish et al., 2006). K_IR_ channels are an exception to this rule (Wu et al., 2007; Kochukov et al., 2014; Sancho et al., 2017; Longden et al., 2017), an observation that raises important questions in the cerebral circulation. Is K_IR_ more than a background conductance that passively sets arterial V_M_? (Nelson and Quayle, 1995; Standen and Quayle, 1998) Could each channel population be distinctively regulated and sensitive to key stimuli including hemodynamic forces? The first step to answer these key questions is to characterize the electrical, molecular and pharmacological profile of cerebral arterial K_IR_ channels (Nelson and Quayle, 1995; Bradley et al., 1999; Schram et al., 2003). Our electrophysiological analysis confirmed the presence of a robust K_IR_ current in smooth muscle and endothelial cells, whose Ba^2+^ sensitivity aligned with K_IR_2.x subunits. A qPCR screen subsequently revealed mRNA for K_IR_2.1/ K_IR_2.2/K_IR_2.4 in both cell types, while Western blot and immunolabeling only detected K_IR_2.1 and K_IR_2.2 at the protein level (Figure 1). These findings align well with previous work (Nelson and Quayle, 1995; Sancho et al., 2017; Schram et al., 2003) and suggest that the K_IR_2.1 and K_IR_2.2 subunits not only dominate expression in the cerebral vasculature but likely form heteromultimeric channels.

### Cerebral Arterial K_IR_2.x Channels are Uniquely Regulated by Membrane Lipids

Vascular K_IR_ channels are often considered poor regulatory targets as stimuli that mobilize classic G-protein coupled receptors routinely fail to alter intrinsic activity (Quayle et al., 1997; Sonkusare et al., 2016). This perspective does not consider recent work, which has identified two membrane lipids as unique modulators of these channels (Huang et al., 1998; Rohacs et al., 2003; Hilgerman, 2007). PIP_2_, a minor component of plasma membranes, has received particular attention given its ability to stabilize K_IR_2.x channels in a preferred open state (Huang et al., 1998; Rohacs et al., 1999; Enkvetchakul et al., 2005). Consistent with this lipid facilitating K_IR_2.x activity, PIP_2_ depletion impaired the endothelial current at voltages where K_IR_ dominates (-100 mV rather than +20 mV). This pattern of endothelial K_IR_ inhibition was repeatable irrespective of how PIP_2_ was depleted, including intracellular ATP modifications, PI kinase blockade, and direct PIP_2_ sequestration (Figures 2 & 3). In contrast, the smooth muscle K_IR_ current was insensitive to PIP_2_ manipulations, a surprising observation but one indicative of each channel pool being distinctly regulated.

Cholesterol is a second membrane lipid able to alter ion channel function in the vascular wall (Romanenko et al., 2002; Fang et al., 2006). For K_IR_, the binding of this lipid through sterol-protein interactions induces a conformational change within the channel pore, promoting its stabilization in a preferred closed state (Romanenko et al., 2004). Respectful of these observations, this study began exploring the modulatory role of cholesterol, with emphasis on smooth muscle K_IR_ as this channel population displayed limited sensitivity to PIP_2_. Findings revealed a pronounced sensitivity of smooth muscle K_IR_ channels to cholesterol, with depletion approaches increasing intrinsic activity by ~100% (Figure 4). In contrast, endothelial K_IR_ channels were markedly less sensitive, with MβCD treatment elevating activity by only ~30%. This differential effect is intriguing and one perhaps indicative of K_IR_ channels localizing to distinct plasma membrane structures. As cholesterol is enriched in caveolae, a subset of experiments next explored whether disrupting Cav1, the major caveolin protein expressed in vasculature, also impacted vascular K_IR_ activity. A blocking peptide directed against Cav1 and introduced via the patch pipette notably increased the smooth muscle K_IR_ current (~40%). A similar effect was absent in endothelial cells, a finding consistent with a greater relative portion of smooth muscle K_IR_ channels residing in caveolae. Building on these observations, immunolabeling techniques (proximity ligation assay) confirmed that K_IR_2.x subunits reside in close association with Cav1 (<40nM) while immuno-electron microscopy structurally placed K_IR_2.1 and K_IR_2.2 within smooth muscle caveolae (Figure 4). These findings cumulatively indicate that each K_IR_ channel population is distinctly regulated, with endothelial and smooth muscle K_IR_ channels predisposed to sitting in an active or silent state, respectively (Rosenhouse-Dantsker et al., 2014; Rosenhouse-Dantsker et al., 2013).

### Lipid-K_IR_2.x Interactions Confer Hemodynamic Sensing in Cerebral Arteries

It is surprising given the rich observational base of model systems (Wu et al., 2007; Ahn et al., 2017; Hoger et al., 2002; Lieu et al., 2004), that native cell studies rarely venture beyond agonists to consider whether hemodynamic forces modulate vascular K_IR_ channels. This study consequently explored K_IR_ channel sensitivity to pressure and flow along with their potential ability to regulate cerebral arterial tone. Using freshly isolated cells, this study showed that stimuli mimicking pressure (hyposmotic challenge/pipette pressurization) alter smooth muscle K_IR_ activity without impacting the same current in endothelial cells. These findings are intriguing as they imply that smooth muscle K_IR_ channels work in concert with pressure-sensitive transient receptor potential channels (Welsh et al., 2002; Earley et al., 2004) to drive myogenic depolarization. This functional perspective aligned with myography observations which noted that Ba^2+^-induced constriction, an index of K_IR_ activity, was greater at low (15 mmHg) rather than high (80 mmHg) intravascular pressure in denuded rat cerebral arteries (Figure 5). With both pressure and cholesterol able to suppress smooth muscle K_IR_ activity, we next considered whether this lipid modulator fostered hemodynamic sensing. Consistent with this, cholesterol depletion prevented pressure stimuli from inhibiting smooth muscle K_IR_ channels (Figure 6). More strikingly, cholesterol depletion in intact arteries facilitated Ba^2+^-induced constriction at 80 mmHg, a finding consistent with this lipid conferring pressure sensitivity to smooth muscle K_IR_ channels.

Flow is a second hemodynamic force likely to affect endothelial ion channels given the positioning of these cells in the arterial wall. While the study of flow sensitivity is logical, the task is challenging as native cells are difficult to harvest and laminar flow chambers must be developed. Regardless, our patch observations are the first to show that laminar flow potentiates the cerebral arterial endothelial K_IR_ current (Figure 7). Akin to the mesentery (Ahn et al., 2017), this potentiation facilitated flow-mediated dilation in cerebral arteries, as intraluminal Ba^2+^ perfusion effectively impaired this hemodynamic response. As both flow and PIP_2_ facilitated endothelial K_IR_ activity, the potential link between this membrane lipid and flow activation was examined. Sequestration of PIP_2_ markedly supressed flow-induced activation of endothelial K_IR_ channels while cholesterol manipulations had no effect (Figure 7). Aligning with patch observations, intraluminal perfusion of neomycin through myogenically active, endothelial intact, vessel segments effectively abolished flow-mediated vasodilation. Together, these findings begin to establish a new paradigm where specific lipid-K_IR_ interactions are essential to hemodynamic sensing and the enabling pressure and flow control in the cerebral vascular tone.

### The Interplay of K_IR_ Channel Pools in the Control of Arterial Tone

It is challenging to picture how ion channel populations, varying in number and distributed among coupled cell populations, work cooperatively in the vessel wall. To address this conceptual issue, we employed computational models, incorporating relevant structural/electrical properties, to quantify the behavior of a single virtual artery (Hald, 2016) or a 3D network of surface vessels and penetrating arterioles (Shih et al., 2009). Alone or as part of a virtual network, each vessel consisted of 1-2 layers of smooth muscle and 1 layer of endothelium. Vascular cells were arranged perpendicular and parallel to the vessel’s long axis and linear resistors representing gap junctions provided interconnectivity. The ionic properties of each cell were modeled as two components, one reflecting K_IR_ and the second representing all other currents. These modeled currents were based on whole cell recordings from smooth muscle and endothelial cells bathed in physiological salt solutions (Figure 8). Our simulations revealed that subtle changes in the smooth muscle or endothelial K_IR_ current induced balanced changes in V_M_/diameter in both computational structures. These balanced responses are important as they indicate that the embedded structural/electrical properties are reasonably representative of native vascular tissue. Equally significant, they reinforce the view that current amplitude, in particular K_IR_, need not be large to have functional impact, as long as cells work in concert and ionic membrane resistivity remains high. In considering smooth muscle K_IR_ channels as a hemodynamic sensor, the impact of other channels in driving myogenic depolarization needs to be considered (Welsh et al., 2000; Welsh et al., 2002). This was illustrated in the channel synergy simulations, where the ability of a pressure-sensitive, non-selective cation current to depolarize vascular cells was noted to increase when K_IR_ channels were simultaneously inhibited.

### Summary

Optimal neural activity relies on balanced blood flow delivery, a process intimately linked to V_M_ and the maintenance of cerebral vascular tone. Arterial V_M_ is set by the sharing of charge and the dynamic interplay among ion channels typically expressed in smooth muscle or the endothelium. Contrary to convention, K_IR_ channels are present in both cell types, an unusual arrangement for an integral membrane protein presumed to operate as a background conductance. Moving beyond this stereotypic view, we re-envisioned K_IR_ as a channel regulated by unique signal transduction molecules that senses the surrounding hemodynamic environment. In detail, our data revealed that K_IR_ channels are distinctively modulated by PIP_2_ and cholesterol depending upon their cellular location; these specific lipid-K_IR_ interactions enable endothelial and smooth muscle K_IR_ channels to respond to flow and pressure, respectively (Figure 9). Computational modeling then showed how the two channel pools worked cooperatively to create an integrated K_IR_ current to set basal V_M_ and tone in the cerebral circulation. The loss of either conductance would logically impact pressure or flow sensing. Indeed, the absence of K_IR_ in mesenteric smooth muscle may explain the modest myogenic response often observed in this vessel bed (Smith et al., 2008). Pathophysiological events that reduce membrane PIP_2_ content could also render the endothelium insensitive to flow regulation, an example being this lipid’s stark drop with advancing Alzheimer’s disease (Arancio, 2008). Alternatively, dyslipidemia where plasma cholesterol rises, could if membrane lipids change accordingly, push endothelial K_IR_ channels to reside in a preferred silent state and insensitive to flow activation (Fancher et al., 2018).

## Methods

### Animal Procedures

The Animal Care Committee at the University of Western Ontario provided protocol approval of all procedures in accordance with the Canadian Council on Animal Care. Briefly, female Sprague-Dawley rats (10-12 weeks of age) were euthanized by carbon dioxide asphyxiation. The brain was carefully removed and placed in cold phosphate-buffered saline solution (PBS; pH 7.4) containing (in mM): 138 NaCl, 3 KCl, 10 Na_2_HPO_4_, 2 NaH_2_PO_4_, 5 glucose, 0.1 CaCl_2_ and 0.1 MgSO_4_. Third-order middle and posterior cerebral arteries were dissected out of surrounding tissue and cut into 2-mm segments.

### Isolation of Cerebral Arterial Smooth Muscle and Endothelial Cells

Smooth muscle cells from cerebral resistance arteries were enzymatically isolated as previously described (Anfigenova et al., 2011). Briefly, arterial segments were placed in an isolation medium (37°C, 10 min) containing (in mM): 60 NaCl, 80 Na-glutamate, 5 KCl, 2 MgCl_2_, 10 glucose and 10 HEPES with 1 mg/mL BSA (pH 7.4). Vessels were then exposed to a two-step digestion process that involved: 1) a 15-min incubation in isolation medium (37°C) containing 0.5 mg/mL papain and 1.5 mg/mL dithioerythritol; and 2) a 10-min incubation in isolation medium containing 100 µM Ca^2+^, 0.7 mg/mL type-F collagenase and 0.4 mg/mL type-H collagenase. Following treatment, tissues were washed repeatedly with ice-cold isolation medium and triturated with a fire-polished pipette. Isolated smooth muscle cells were identified by their characteristic spindle-like shape and contractile behaviour. Liberated cells were stored in ice-cold isolation medium for use the same day within ~ 5 hours.

Endothelial cells were isolated as described previously (Sancho et al., 2017). Briefly, arterial segments were placed in an isolation medium (37°C, 10 min) containing (in mM): 140 NaCl, 5.5 KCl, 1 MgCl_2_, 1.2 NaH_2_PO_4_, 5 glucose, 2 Na^+^ pyruvate, 0.02 EDTA and 10 HEPES with 0.1 mg/mL BSA (pH 7.4). Vessels were then exposed to a two-step digestion process that involved the following: 1) a 30-min incubation in isolation medium (37°C) containing 1mg/mL BSA, 100 µM Ca^2+^, 1mg/mL papain and 1 mg/mL dithioerythritol; and 2) a 10-min incubation in isolation medium containing 1 mg/mL BSA, 100 µM Ca^2+^, 0.9 mg/mL type-F collagenase, 0.6 mg/mL type-H collagenase, 5 mg/mL elastase and 1 mg/mL trypsin inhibitor. Tissues were then washed repetitively with ice-cold isolation medium and gently triturated with a fire-polished pipette. This isolation procedure yielded small groups of endothelial cells and they were clearly identifiable by their rough shape and lack of voltage dependent K^+^ conductances. Isolated cells were stored in ice-cold isolation medium and used the same day for up to 4 hours.

### Electrophysiological Recordings

Conventional patch-clamp electrophysiology was used to measure whole-cell currents in both isolated smooth muscle and endothelial cells. Briefly, recording electrodes (resistance of 5-8 MΩ when filled with solution) were pulled from borosilicate glass microcapillary tubes (Sutter Instruments, Novato, CA), covered in dental wax to reduce capacitance, and backfilled with pipette solution containing (in mM): 5 NaCl, 35 KCl, 100 K-gluconate, 1 CaCl_2_, 0.5 MgCl_2_, 10 HEPES, 10 EGTA, 2.5 Na_2_-ATP and 0.2 GTP (pH 7.2). To attain whole-cell configuration, a pipette was then gently lowered on to a cell and negative pressure applied to achieve a giga-ohm seal and rupture the membrane. Cells were voltage-clamped at a holding membrane potential of -60 mV and equilibrated for 15 min in a bath solution containing (in mM): 135 NaCl, 5 KCl, 0.1 MgCl_2_, 10 HEPES, 10 glucose and 0.1 CaCl_2_ (pH 7.4). Whole-cell currents were recorded on an Axopatch 200B amplifier (Axon Instruments, Union City, CA, USA), filtered at 1 kHz, digitized at 5 kHz and stored on a computer for subsequent offline analysis with Clampfit 10.3 software (Molecular Devices, Sunnyvale, CA). Cell capacitance ranged between 14-18 pF in smooth muscle cells and 4-8 pF in endothelial cells was measured with the cancelation circuity in the voltage-clamp amplifier. Cells that displayed a noticeable shift in capacitance (>0.3 pF) during experiments were excluded for analysis. A NaCl-agar salt bridge between the Ag-AgCl reference electrode and the bath solution was used to minimize offset potentials (<2 mV). All experiments were performed at room temperature (~22°C).

Ba^2+^-sensitive K_IR_ currents were quantified by elevating extracellular [K^+^] from 5 to 60 mM K^+^ via equimolar replacement of NaCl by KCl. Voltage was then stepped to -100 mV for 100 ms and then ramped to +20 mV at a rate of 0.04 mV/ms. The currents from three trials (5s between trails) were subsequently averaged. To assess Ba^2+^ sensitivity, the preceding protocol was applied to cells, bathed in a 60 mM K^+^ solution while the concentration (1-100 µM) of this K_IR_ inhibitor was sequentially elevated. In a subset of experiments, extracellular [K^+^] was maintained at 60 mM, and different agents that alter PIP_2_ or cholesterol membrane content were added to the pipette solution including: 1) ATP (0, 2.5 or 5 mM); 2) neomycin (50 µM); 3) PIP_2_ (50 µM); 4) wortmannin (50 µM) or adenosine (2 mM); 5) methyl-beta-cyclodextrin ± cholesterol (5 mM); and 6) Cav1 scaffolding domain peptide (Cav1 SDP, 10 µM, corresponding to Cav1 amino acids 82-101) or Cav1 SDP scrambled control (10 µM). In all subsequent experiments, extracellular [K^+^] was sustained at 20 mM and the magnitude of the inward current measured prior to and following diverse mechanical perturbations including: 1) simulated pressure by hyposmotic challenge (reducing bath osmolarity from 300 to 205 mosM by adding/removing 100 mM D-mannitol from the bath solution) (Wu et al., 2007) or negative pipette pressurization (-15 to -45 mmHg using a through a pneumatic transducer (DPM-1B, BioTek Instruments, VT, USA); and 2) laminar flow through a custom patch-clamp chamber. Shear stress was calculated to ~10 mPa, a value that sustains patch seal integrity.

### qPCR Analysis

Total RNA was extracted from middle/posterior cerebral arteries (~2), smooth muscle (~200) and endothelial cells (~100) using the RNeasy plus micro kit (Qiagen) according to instructions. Reverse transcription was performed using the Quantinova reverse transcription kit (QIAGEN). For the negative control groups, all components except the reverse transcriptase were included in the reaction mixtures. qPCR was performed using Primer Time qPCR primers (Table S2) and the Kapa SYBR Fast Universal qPCR Kit (Kapa Biosystems). Rat β-actin (ACTB) was utilized as the reference gene. Control reactions and those containing cDNA from cerebral arteries were performed with 1 ng of template per reaction. Due to the very small quantities of RNA obtained from isolated smooth muscle and endothelial cells, the entire cDNA yield from each preparation was used to assay the full set of test and housekeeping genes. The running protocol included 45 cycles consisting of 95°C for 5 s, 55°C for 15 s and 72°C for 10 s using an Eppendorf Realplex 4 Mastercycler. PCR specificity was checked by dissociation curve analysis. Assay validation was confirmed by testing serial dilutions of pooled template cDNAs suggesting a linear dynamic range of 0.1-100 ng template and yielded percent efficiencies ranging from 85-108%. Template controls did not yield detectable fluorescence. Expression of the K_IR_2.x isoforms in cerebral arteries, endothelial or smooth muscle cells relative to control tissue was determined using the relative expression software tool (REST) version 2.0.13 (Pfaffl et al., 2002).

### Western Blotting

Intact cerebral arteries were placed in 100 µl ice-cold lysis buffer (pH 7.4) containing (in mM) 20 Tris-HCl, 100 NaCl, 10 MgCl2, 1 EDTA, 1 EGTA, 2 sodium pyrophosphate, and 1 beta-glycerophosphate with 0.5% Nonidet P40 and a cocktail of mammalian protease inhibitors (200 mM Na_3_VO_4_, 0.5 M NaF, 1mg/mL aprotinin, 1mg/mL leupeptin, 100 mM PMSF, and 100 mM DTT). Samples were mechanically disrupted and centrifuged (1.5 min, 13,000 rpm). Supernatant containing protein was then transferred to a clean tube, assayed for total protein using Pierce BCA Protein Assay Kit, and stored at -20°C for up to 1 wk. Following protein quantification, 10 µg of protein were loaded to run on a 4-10% gradient SDS-PAGE gel (80 min, 100 V). Protein was transferred to a nitrocellulose membrane, blocked for 1 hour (5% non-fat dairy milk in Tris-buffered saline) and incubated (2h, 20-22°C) with a primary antibody (K_IR_2.1, K_IR_2.2, or K_IR_2.4 see Table S3) diluted in 1% milk-Tris buffer. The membrane was subsequently washed and incubated with goat polyclonal anti-rabbit horseradish peroxidase-conjugated secondary antibody (1:10,000) diluted in 1% milk-Tris buffer (1h, 20-22°C). Washing was repeated, and the blot was developed using Amersham ECL Prime Western Blotting Reagent and then imaged on a Gel Doc using Image Lab 3.0 software (Bio-Rad).

### Immunohistochemistry

K_IR_ 2.1, K_IR_ 2.2, K_IR_ 2.4 and caveolin-1 (Cav1) protein expression was assessed in isolated cerebral arterial smooth muscle cells or in endothelial layers from whole-mount cerebral artery preparations. Briefly, isolated cells were air dried onto microscope cover glass, and longitudinally opened vessels were mounted onto a Sylgard-block, and both were then fixed/ permeabilized in PBS (pH 7.4) containing 4% paraformaldehyde and 0.2% Tween 20. Fixed cells/arteries were blocked/permeabilized (1h, 22°C) with a quench solution (PBS supplemented with 3% donkey serum, and 0.2% Tween 20) and subsequently incubated overnight (4°C, humidified chamber) with primary antibodies: anti-K_IR_2.1, anti-K_IR_2.2, anti-K_IR_2.4, and anti-Cav1 (Table S3) diluted in quench solution. On the next morning, cells/vessels were washed three times in PBS-0.2% Tween 20 and then incubated (1h, 22°C) in a PBS-0.2% Tween 20 buffer containing Alexa Fluor 488 donkey anti-rabbit IgG-secondary antibody (1:200). After further washing, isolated cells and whole-mount preparations were mounted with Prolong Diamond Antifade Mountant with DAPI. Immunofluorescence was detected through a x63 oil immersion lens coupled to a Leica-TCS SP8 confocal microscope equipped with the appropriated filter sets. Smooth muscle cells isolated from mesenteric arteries were used as K_IR_2.x negative controls. Secondary antibody controls were also performed and were negative for nonselective labeling.

### Proximity Ligation Assay (PLA)

The Duolink *in situ* PLA detection kit was employed using freshly isolated smooth muscle cells. Briefly, cells were fixed in PBS containing 4% paraformaldehyde (15 min), and then incubated with PBS-0.2% Tween 20 buffer (15 min) for permeabilization. Cells were then washed with PBS, blocked by Duolink blocking solution (1h, 37°C) and incubated overnight (4°C) with primary antibodies: anti-K_IR_2.1, anti-K_IR_2.2, and anti-Cav1 (Table S3) in Duolink antibody diluent solution. Control experiments employed no primary antibody or only one primary antibody. Cells were labelled with Duolink PLA PLUS and MINUS probes for 1h (37°C). The secondary antibodies of PLA PLUS and MINUS probes are attached to synthetic oligonucleotides that hybridize if present in close proximity (<40 nm). The hybridized oligonucleotides are then ligated and subjected to amplification. The amplified products extending from the oligonucleotide arm of the PLA probe were detected using far red fluorescent fluorophore-tagged, complementary oligonucleotide sequences and a Leica TCS SP8 confocal microscope. Nuclei were stained with DAPI.

### Immunogold Labelling

Animals were anaesthetized with sodium pentathol (100 mg/kg; ip) and perfusion fixed in 0.2% glutaraldehyde and 2% paraformaldehyde in 0.01 M PBS (pH 7.4). Cerebral artery segments (~2 mm in length) were rinsed (3 x 5 min) and stored in PBS before processing in a Leica EMPACT 2 high-pressure freezer using 0.7% low melting agarose as a cryoprotectant. Samples were then freeze-substituted in a Leica AFS2 into 0.2% uranyl acetate in 95% acetone (from -85 to -50°C) and infiltrated with Lowicryl (at -50°C; ProSciTech) before UV polymerisation (2 d each at -50 and 20°C).

Individual serial thin transverse sections (~100 nm) were mounted on Formvar-coated slot grids and processed for antigen localization as for confocal immunohistochemistry (see Table S3 for antibody details), although the secondary used was 5 or 10 nm colloidal gold-conjugated antibody (1:40; 2 hours) in 0.01% Tween-20. Controls included peptide block (where available) and omission of the primary antibody. Sections were imaged at x40-60,000 on a JEOL transmission electron microscope at 16 MP (Emsis, Morada G3). Images were uniformly adjusted for brightness and contrast in Photoshop CC.

### Vessel myography

Cerebral arterial segments were mounted in a customized arteriograph and superfused with warm (37° C) physiological salt solution (PSS; pH 7.4) containing (mM): 119 NaCl, 4.7 KCl, 20 NaHCO_3_, 1.7 KH_2_PO_4_, 1.2 MgSO_4_, 1.6 CaCl_2_, and 10 glucose. Arteries were equilibrated for 30 min (15 mmHg intravascular pressure) and contractile responsiveness assessed by brief (~ 10 s) exposure to 55 mM KCl. In a subset of experiments, pressure-induced responses were studied in endothelium-denuded arteries by passing an air bubble through the vessel lumen; successful removal was confirmed by the loss of bradykinin-induced dilations. Following equilibration, cerebral arteries were maintained at low (15 mmHg) or high (80 mmHg) intravascular pressure in the absence or presence of BaCl_2_ (30 µM; 2-3 min) ± methyl-beta-cyclodextrin (5 mM, 30 min pretreatment). Third-order branches of rat mesenteric artery were used as negative control for smooth muscle K_IR_.

In all subsequent experiments, flow-induced responses were studied in endothelium-intact arteries mounted in a pressure-flow chamber where the inflow and outflow pressures were controlled by a pressure servo-control system (Living Systems Instrumentation, USA). The bore of glass pipettes was matched to each other to achieve equal resistance. Following equilibration, intravascular pressure were slowly raised from 15 mmHg to 60 mmHg and cerebral arteries were then exposed to bradykinin (15 µM) to confirm the presence of intact endothelium. Vascular reactivity was then monitored prior to and following stepwise increases in intraluminal flow (2, 4, and 6 µl/min; 5 min steps) using a flow pump. Changes in diameter were measured under control conditions or in the presence of external/intraluminal BaCl2 (30 µM) or intraluminal neomycin (50 µM). Arterial diameter was monitored by using a x10 objective and an automated edge detection system (IonOptix, USA).

### Computational Modeling

A 3D model of an arterial network was constructed using a generative modeling approach (Hald, 2016). Briefly, a single virtual vessel of defined length and diameter comprised of one endothelial cell layer with endothelial cells (EC) oriented parallel to the vessel axis and one or more smooth muscle cell layers with smooth muscle cells (SMC) oriented perpendicular to the vessel axis. Each layer represent a 3D space filled with cells of a predefined volume that communicate to all neighboring cells via homocellular and myoendothelial gap junctions (vessel ends being electrically sealed).

In order to create a 3D network, coordinates of vessel bifurcations and network ends were obtained from Shih et al 2009 (Shih et al., 2009). Assuming vessels to be straight cylinders, vessel lengths and connectivity were calculated (meandering vessels were approximated using piecewise linear curve fits) (Knot and Nelson, 1998). Each surface vessel was covered with two smooth muscle layers and was assumed to have a resting diameter of 75 µm, whereas penetrating arterioles were 400 µm long, covered with a single layer of smooth muscle and assumed to have a resting diameter of 30µm. At each bifurcation, endothelial cells at the end of an incoming branch were divided into two sets; the number of ECs in each set being dependent on the ratio of ECs in the two opposing branches. Endothelial cells within each set were coupled to endothelial cells from a single opposing branch while smooth muscle cells, due to their circumferential orientation, were coupled to at least one smooth muscle cell from each opposing branch, at the bifurcation (Hald, 2016).

In each cell, electrical dynamics were described using classical Hodgkin-Huxley formalism:

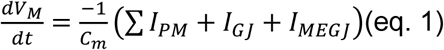

where*C_m_*is cell capacitance, *I_PM_* is current across the plasma membrane (PM),*I_GJ_*is current through homocellular gap junctions (GJ) and *I_MEGJ_* is current through myoendothelial gap junctions (MEGJ), i.e. between the EC layer and the innermost SMC layer. Ohmic relations *Ij* · *R_j_* = *ΔV_j_*, where *ΔV_j_* is the transjunctional potential, described both GJ and MEGJ currents. *I_PM_* was modeled via simple polynomial fits to whole-cell mode patch-clamp recordings (steady state IV-curves) of isolated endothelial and smooth muscle cells in physiological solution.

Reversal potentials of patched cells may not adhere to resting potentials of cells embedded in intact tissue because of e.g. partial cell degradation during enzymatic isolation, temperature differences, and/or effects of cell dialysis due to artificial patch solutions. The polynomials were aligned to the physiologically relevant resting potential of -45mV either by applying weighted fitting (EC fit) or by addition of a small linear “correcting current” (SMC fit). The resulting polynomial fits were described by:

I_PM,EC_ (V_M_) = 0.276 + 7.58 · 10^−3^ V_M_ – 1.59 · 10^−4^ V_M_^2^ – 7.82 · 10^−6^ V_M_^3^ – 7.90 · 10^−8^ V_M_^4^

I_PM,SMC_(V_M_) = 1.14 + 0.08 V_M_ + 2.18 ·10^−3^ V_M_ ^2^ + 2.92 ·10^−5^ V_M_ ^3^ + 1.92 ·10^−7^ V_M_ ^4^ + 5.58 ·10^−10^ V_M_ ^5^

Similarly, polynomials were fitted to K_IR_-channel currents that in turn could be multiplied with a certain factor and added to or subtracted from the plasma membrane current. The resulting polynomial fits were described by:

I_KIR,SMC_ (V_M_) = -0.092 + 4.21 · 10^−3^ V_M_ + 1.04 · 10^−3^ V_M_^2^ + 3.24 · 10^−5^ V_M_^3^ + 3.85 · 10^−7^ V_M_ ^4^ + 1.62 · 10^−9^ V_M_^5^

I_KIR,EC_(V_M_) = -0.124 + 1.38 V_M_ · 10^−3^ V_M_ + 4.34 ·10^−5^ V_M_^2^ − 7.19 ·10^−6^ V_M_^3^ − 8.64 ·10^−8^ V_M_ ^4^

The resulting partial differential equation (PDE) was reduced to a set of ordinary differential equations (ODE) using the finite difference method and solved using CVODE 2.7.0 from Sundials (Hindmarsh et al., 2005) using relative tolerance of 1E-4 (absolute tolerance of 1E-9).

Following simulation of V_M_ within each cell of the vascular network, the electrical response was translated into a change in inner diameter using a sigmoid function:

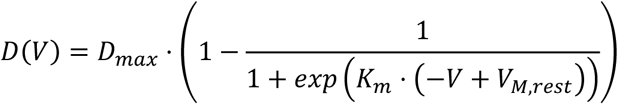

where V_M,rest_ is the resting potential that corresponds to the electrical model. For penetrating arterioles, D_max_ = 50 µm and for surface vessels D_max_ = 120 µm.

To estimate K_m_, a constant determining the dynamic range of excitation-contraction coupling, a minimal diameter, D_min_, was added to the sigmoid and this function was fitted to the observed relationship between simultaneously measured V_M_ and outer vessel diameter in cerebral vessels (see Figure 8-figure supplement 1) (Knot and Nelson, 1998); from the fit, K_m_ = 0.134.

*Limitations:* The computational models are simple which allows for direct estimation of model parameters from experimental data but also leaves out a number of important regulators of arteriolar function. In particular, the steady state model of plasma membrane currents disregard the impact of paracrine/endocrine factors from nervous or immune origin, and disregards dynamics of individual ion channels. Thus, addition/subtraction of individual currents to the overall plasma membrane current does not provide an exact or complete representation of activation/inactivation as other ion channels may likely respond in non-linear fashion.

### Solutions and Chemicals

All buffers, chemical and reagents originated from Sigma-Aldrich unless otherwise stated. Papain was acquired from Worthington; elastase, Cav1 SDP and Cav1 SDP scrambled control from Calbiochem and DiC8-PIP_2_ from Echelon. Primary antibodies against K_IR_2.1, K_IR_2.2 were obtained from Alomone whereas those directed against K_IR_2.4 and Cav1 were purchased from Thermofisher Scientific. Alexa Fluor 488 donkey anti-rabbit IgG-secondary antibody, Prolong Diamond Antifade Mountant and Pierce BCA Protein Assay Kit were also obtained from Thermofisher Scientific. Peroxidase goat anti-rabbit secondary antibody was purchased from Jackson Laboratories, and Amersham ECL Prime Western Blotting Reagent was obtained from GE Healthcare.

### Statistical Analysis

Data are expressed as means±S.E; and n indicates the number of cells or arteries or animals. No more than two different experiments were performed on vessels from a given animal. Power analysis was done to calculate sample size sufficient for obtaining statistical significance and n numbers (n=6) reported in previous publications were considered, in experiments involving cells, vessels and animals. Where appropriate, paired, unpaired *t*-tests, or Repeated Measures (RM) - ANOVA followed by post-hoc Tukey test were performed to compare the effects of a given condition/treatment on whole cell current or arterial diameter. \**P* values ≤ 0.05 were considered statistically significant. Experimental design and data presentation are reported in compliance with the ARRIVE guidelines.

## Acknowledgements

The authors would like to acknowledge the excellent technical support of Suzanne Brett (University of Western Ontario), Michelle Kim and William Gao for assistance with Western blot, Dr. Chris Ellis for his conceptual contributions and Dr. Frank Visser (Hotchkiss Brain Institute, University of Calgary) for assistance with qPCR. Dan Shelley (University of the Sunshine Coast), and Rick Webb and Kathryn Green (University of Queensland) assisted with the Electron Microscopy. This work was supported by operating grants to DGW from the Heart and Stroke Foundation of Canada and Canadian Institute of Health Research. DGW is the Rorabeck Chair in Neuroscience and Vascular Biology at the University of Western Ontario. SLS was supported by Brain Foundation of Australia.

---

**Figure 3-figure supplement 1.**
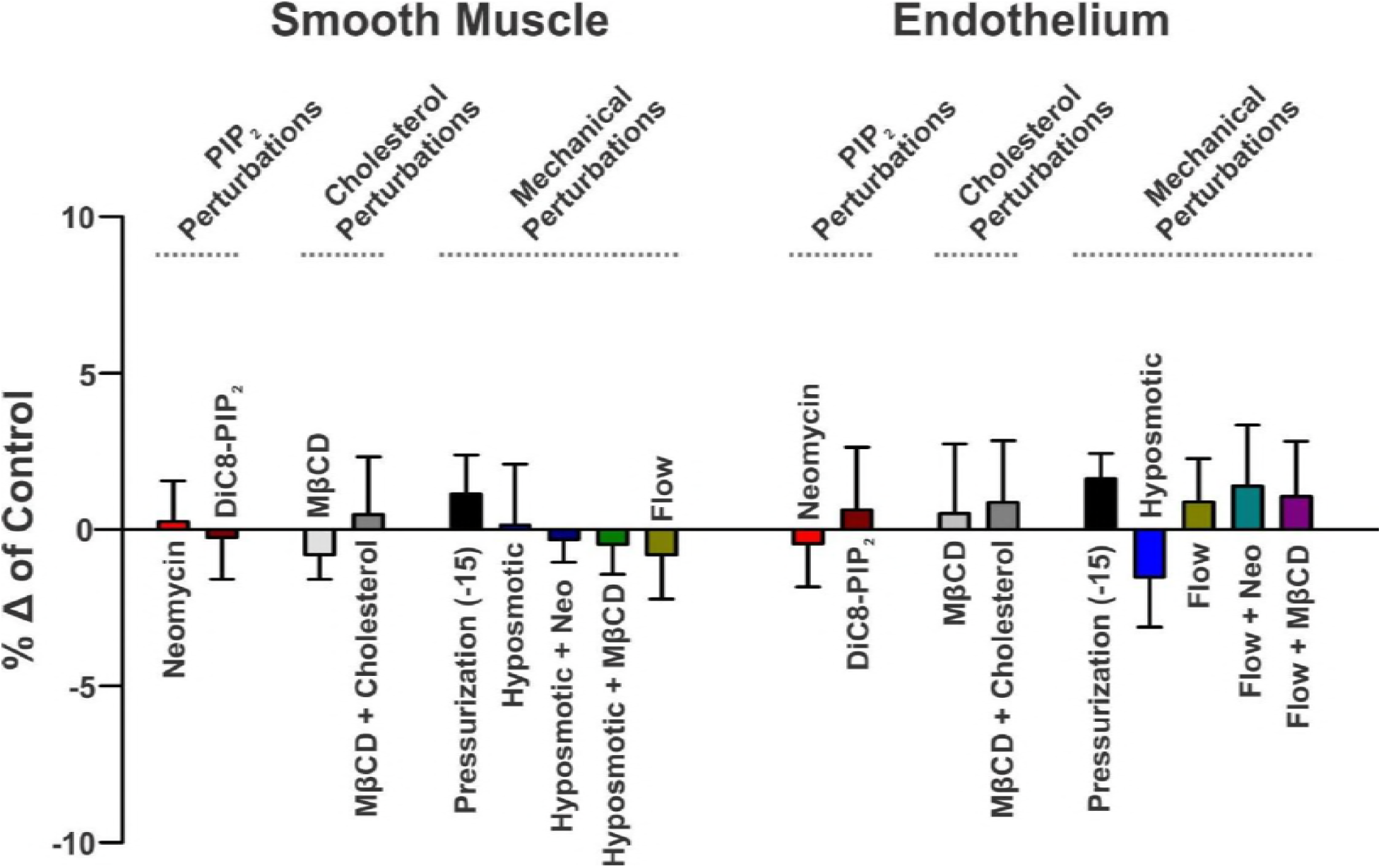
Membrane lipid (PIP_2_ and cholesterol) and mechanical perturbations don’t impact outward currents. Whole-cell currents were measured (voltage ramp, -100 to +20 mV) in smooth muscle (left) and endothelial cells (right) bathed in a 60 (lipid regulation) or 20 (mechanical regulation) mM K^+^ solution. Summary data (n=6-8, unpaired *t* test) of peak outward current (+20 mV) expressed as percent change over the 12 min period. Perturbations were to: 1) PIP_2_ content (50 µM neomycin; 50 µM DiC8-PIP_2_); 2) cholesterol content (5 mM methyl-β-cyclodextrin (MβCD) ± 5 mM cholesterol; and 3) the mechanical state of the cell (pipette pressurization; hyposmotic challenge ± neomycin or MβCD; flow ± neomycin or MβCD). Data are means ± SE.

**Figure 3-figure supplement 2.**
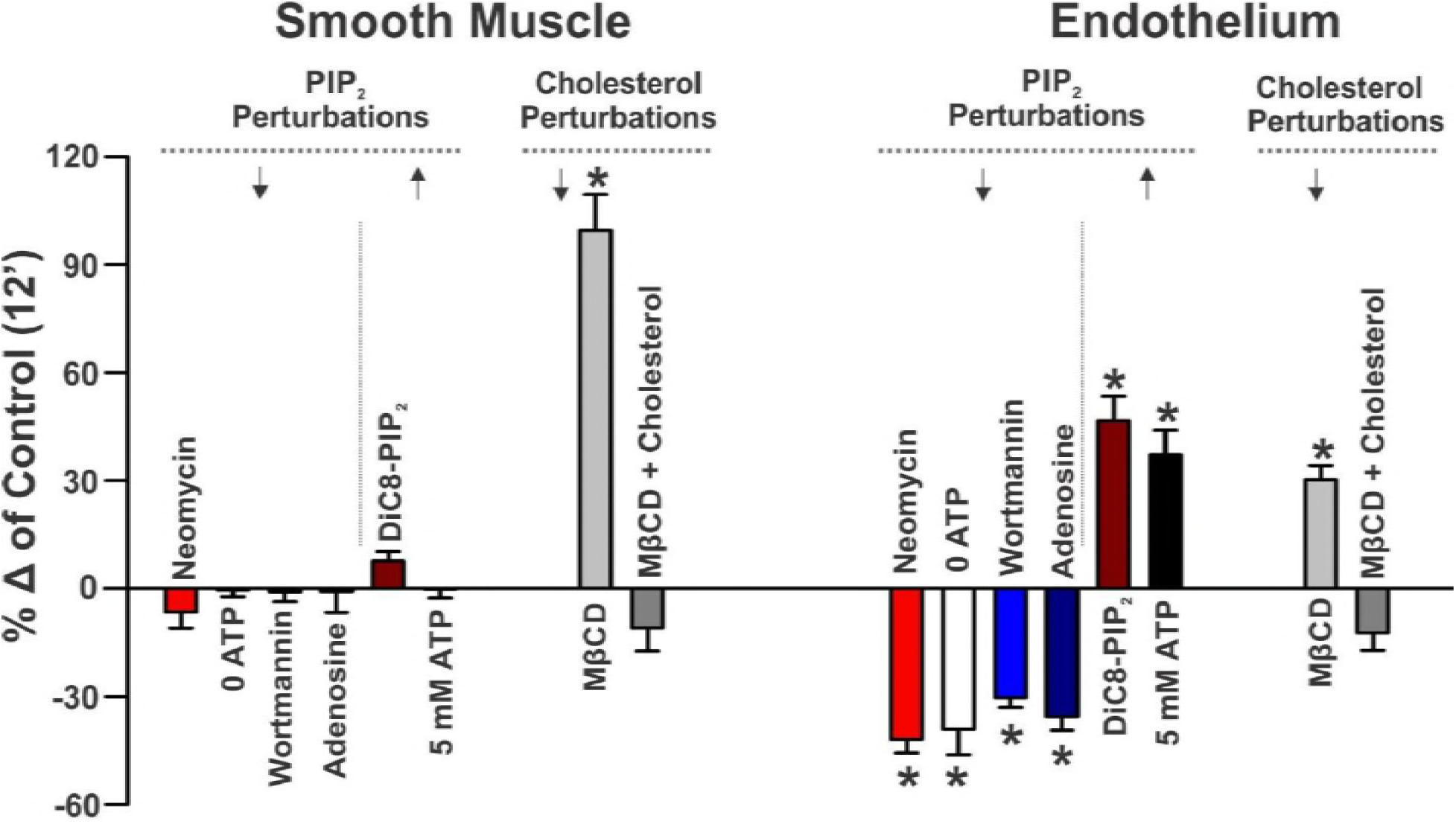
PIP_2_ and cholesterol are the dominant regulators of endothelial and smooth muscle K_IR_ channels, respectively. Whole-cell currents were measured (voltage ramp, -100 to +20 mV) in smooth muscle (left) and endothelial cells (right) bathed in a 60 mM K^+^ solution. Summary data (n=6-8, unpaired *t* test) of peak inward current (- 100 mV) was expressed as percent change over the 12 min recording period. Perturbations were to: 1) PIP_2_ content (50 µM neomycin; 0 or 5 mM ATP; 50 µM wortmannin; 2 mM adenosine; 50 µM DiC8-PIP_2_); and 2) cholesterol content (5 mM methyl-β-cyclodextrin (MβCD) ± 5 mM cholesterol). Data are means ± SE. * denotes significant difference from 0 min.

**Figure 5-figure supplement 1.**
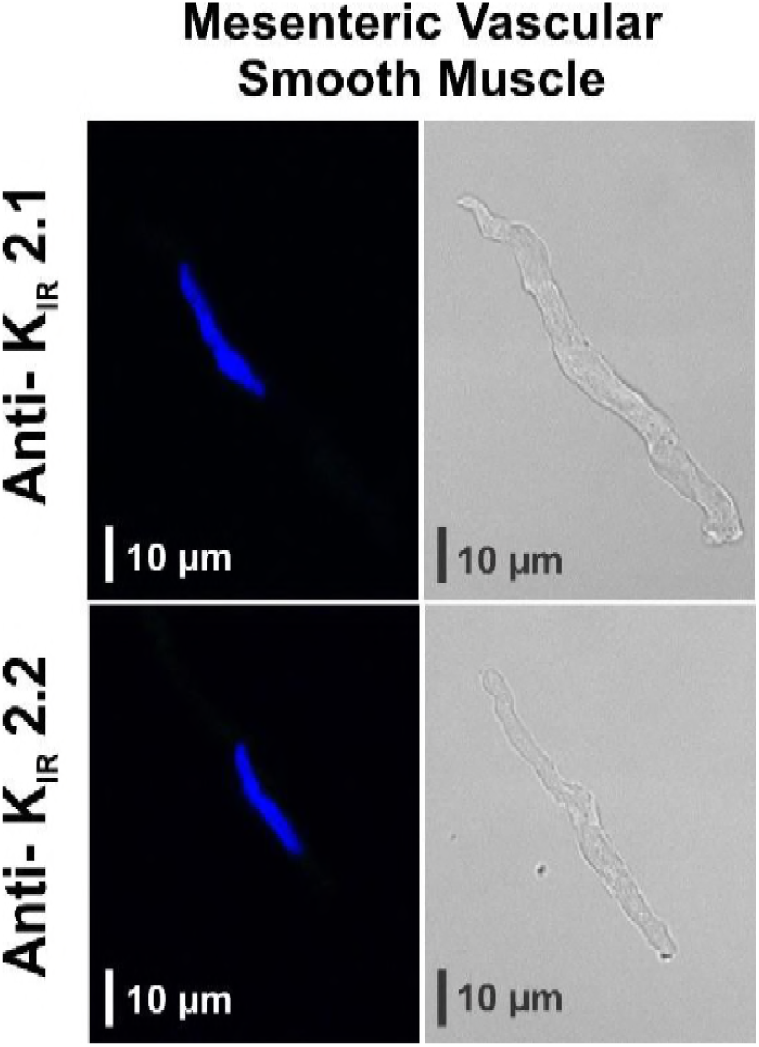
K_IR_2.x channels are absent in native mesenteric arterial smooth muscle cells. Immunohistochemistry of freshly isolated smooth muscle cells (top) reveals no K_IR_ 2.1 or K_IR_2.2 (green) protein labeling. Nuclei were stained with DAPI (blue); assay controls were performed with no primary antibodies. Each experiment was tested on cells for 4 different animals; photomicrographs are representative of 10-20 cells per group.

**Figure 6-figure supplement 1.**
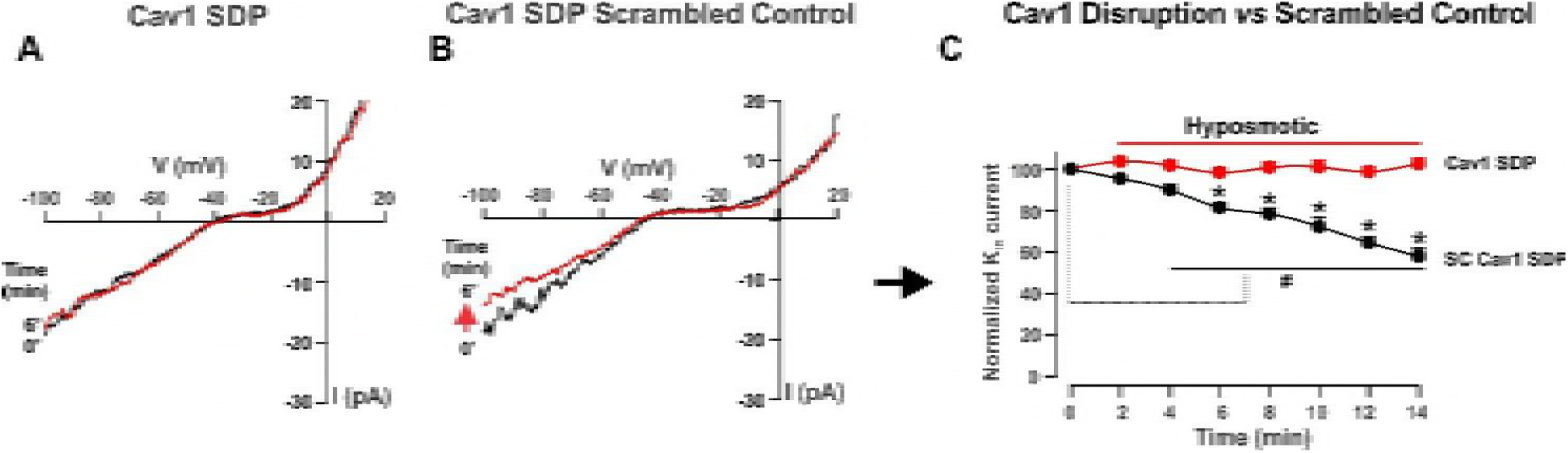
Caveolin-1 disruption impairs swelling-mediated inhibition of smooth muscle K_IR_ channels. Representative traces (**A**-**B**) and summary data (**C;** n=6 cells per group) of the smooth muscle K_IR_ current, prior to and during a hyposmotic challenge where cells were dialyzed with pipette solution containing Cav1 SDP (10 µM), or Cav1 SDP scrambled control (10 µM). RM-ANOVA and Tukey post hoc analysis compared time-course differences in the same cell; unpaired *t* test compared inward current (-100 mV) changes among treatments. Data are means ± SE. ^*, #^ denotes significant difference at *P* ≤ 0.05.

**Figure 7-figure supplement 1.**
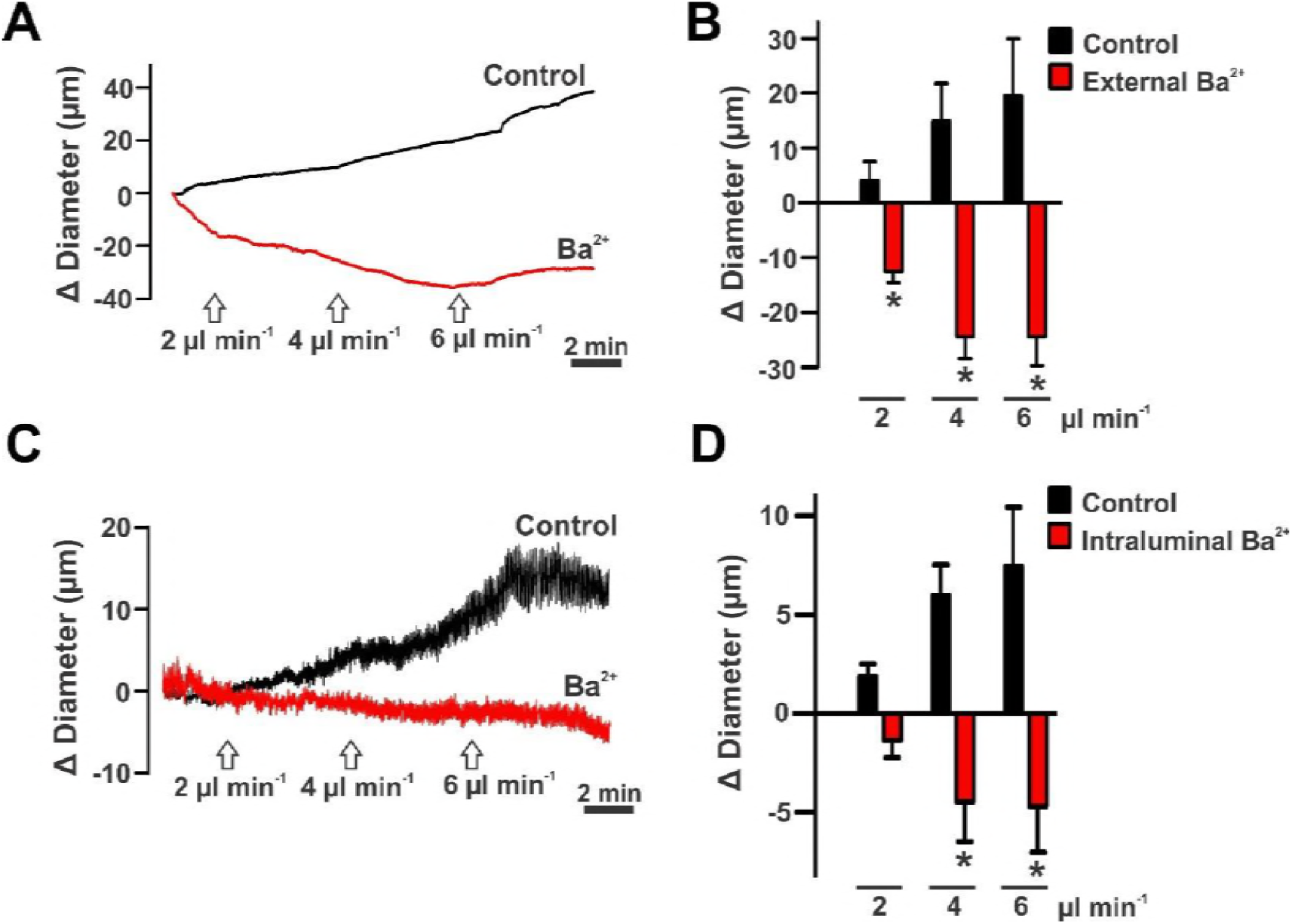
Endothelial K_IR_ channels mediate flow-induced dilation in endothelial intact cerebral arteries. Representative traces and summary plots (n=6 per paired group) comparing the effect of external (A, B) *versus* intraluminal (C, D) Ba^2+^ (30 µM) on flow-mediated responses (0 to 4 or 6 µl/min). Data are means ± SE and * denotes significant difference to control at *P* ≤ 0.05; paired *t* test.

**Figure 8-figure supplement 1.**
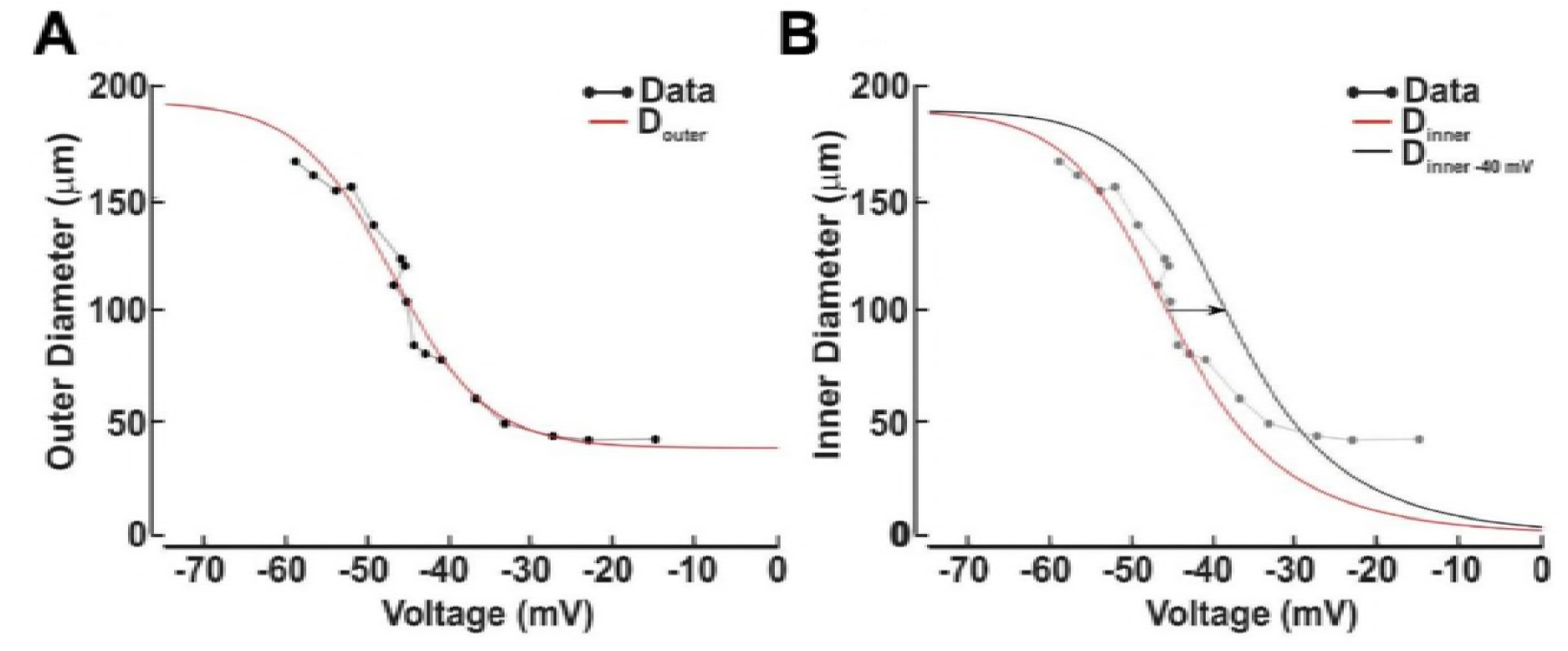
Calculated relationship between diameter and arterial V_M_. **A**, The relationship between arterial V_M_ and outer cerebral arterial diameter was plotted and a sigmoid function (red; see Eq. 1) applied. **B**, Inner diameter was represented by assuming that a vessel is fully contracted at 0 mV and that the cross-sectional vessel area is constant across measurements (red curve). This curve was right-shifted to correspond to a resting V_M_ of -40 mV; black curve).

